# Divisive attenuation based on noisy sensorimotor predictions accounts for excess variability in self-touch

**DOI:** 10.1101/2024.06.20.599826

**Authors:** Nicola Valè, Ivan Tomić, Zahara Gironés, Daniel M Wolpert, Konstantina Kilteni, Paul M Bays

## Abstract

When one part of the body exerts force on another, the resulting tactile sensation is perceived as weaker than when the same force is applied by an external agent. This phenomenon has been studied using a force matching task, in which observers are first exposed to an external force on a passive finger and then instructed to reproduce the sensation by directly pressing on the passive finger with a finger of the other hand: healthy participants consistently exceed the original force level. However, this exaggeration of the target force is not observed if the observer generates the matching force indirectly, by adjusting a joystick or slider that controls the force output of a motor. Here we present the first detailed computational account of the processes leading to exaggeration of target forces in the force matching task, incorporating attenuation of sensory signals based on motor predictions. The model elucidates previously unappreciated contributions of multiple sources of noise, including memory noise, in determining matching force output, and shows that quantifying attenuation as the discrepancy between direct and indirect self-generated forces isolates its predictive component. Our computational account makes the prediction that attenuated sensations will display greater trial-to-trial variability than unattenuated ones, because they incorporate additional noise from motor prediction. Quantitative model fitting of new and existing force matching data confirmed the prediction of excess variability in self-generated forces and provided evidence for a divisive rather than subtractive mechanism of attenuation, while highlighting its predictive nature.

**NEW & NOTEWORTHY:** We formulate a detailed computational account of sensory attenuation in force matching tasks that disambiguates contributions of perceptual, memory and prediction noise in order to isolate a pure measure of attenuation strength. Analysis of data from nearly 500 participants shows that attenuated sensations display increased trial-to-trial variability, consistent with incorporating additional noise inherent to motor prediction. These results support a divisive – rather than subtractive – reduction in the sensation of self-generated forces based on predicted reafference.

## INTRODUCTION

The same tactile stimulus is perceived differently when it is the result of your own action compared to when it has an external source. This phenomenon is often referred to as sensory attenuation, as a self-generated stimulus is typically perceived as weaker than an externally generated one. A predictive mechanism has been proposed for this phenomenon [1–6], inspired by neurophysiological mechanisms of sensory cancellation [7–10]. According to the predictive account, when the motor system generates a voluntary movement, a duplicate of the motor command (an efference copy) is provided to a forward model that integrates it with an estimate of the state of the body to predict sensory consequences of the movement [11]. This prediction is used to attenuate incoming tactile signals (reafference) that are a consequence of the action.

Shergill et al. [2] described a force matching task, in which participants were asked to reproduce the tactile sensation of a target force, delivered to their passive left index finger by a lever attached to a torque motor (Fig. 1A). The reproduction was performed in two conditions: participants either applied a force to the left index finger by pressing on the lever with their right index finger (direct condition, Fig. 1B) or they used their right index finger to adjust the setting of a response device that controlled the force output of the torque motor (indirect condition, Fig. 1C). In both conditions, the participants were asked to modulate the force on their passive left finger until it matched the tactile sensation of the preceding target force.

**Figure 1:**
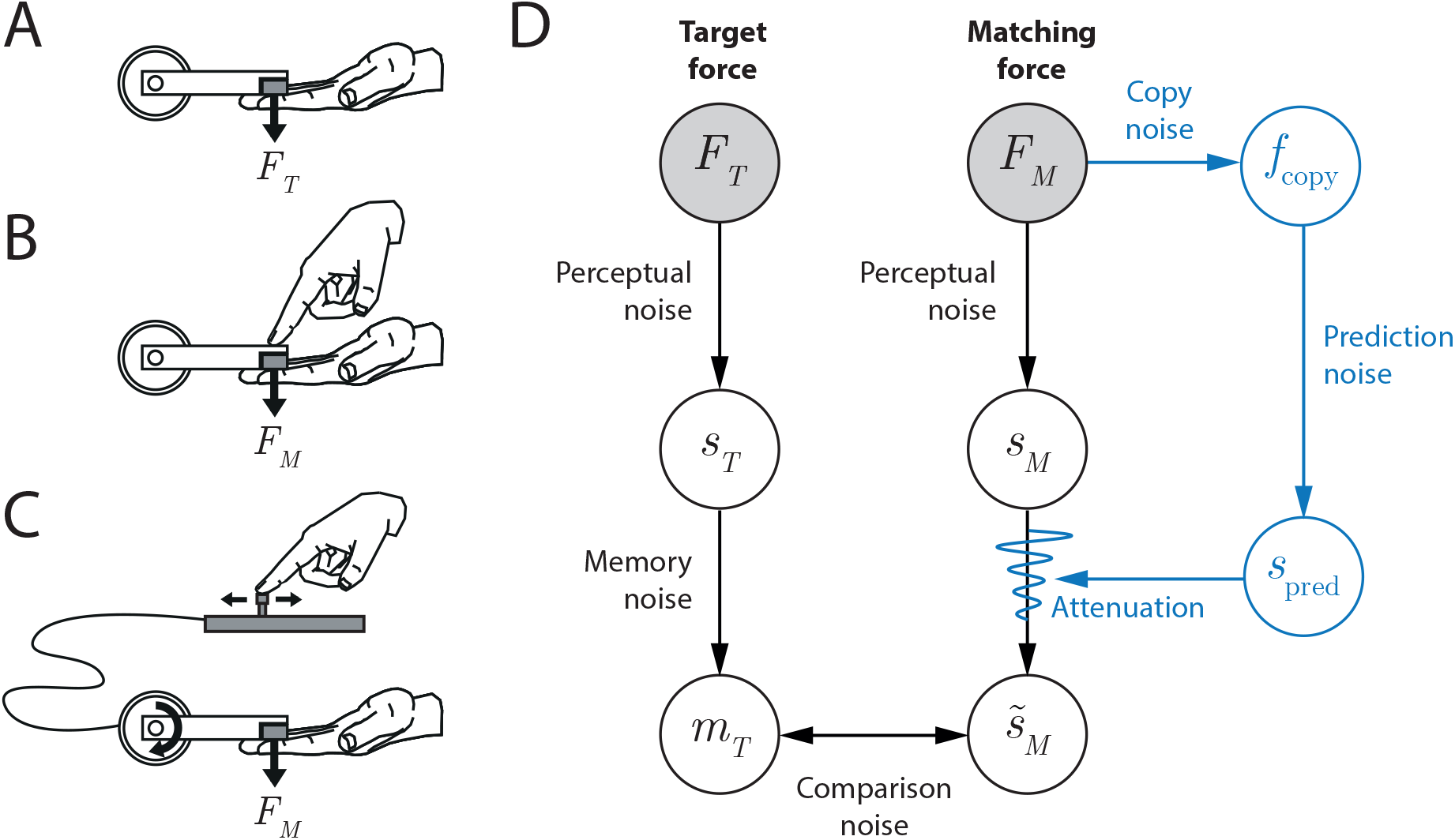
(A–C) The force matching task. (A) A lever attached to a torque motor delivers a target force (*F*_*T*_) to a participant’s passive index finger. (B) In the direct condition, the participant reproduces the force sensation (with force *F*_*M*_) by pressing with the other hand. (C) In the indirect condition, the participant reproduces the force sensation by controlling the torque motor with a slider (shown) or joystick. (D) Diagram of the processes involved in the force matching task. Black: processes involved in both the direct and indirect condition. Blue: model of the predictive mechanism leading to attenuation. This prediction is effective only in the direct condition. See the text for a more extensive description.

The results showed that healthy participants produced matching forces that consistently exceeded the target forces in the direct condition, whereas matching forces were much closer to veridical in the indirect condition. Since the manner of presentation of the target force was identical in both conditions, the authors concluded that the exaggerated matching forces in the direct condition were caused by a relative attenuation of the tactile sensation induced in the passive finger by the participant’s own action. Moreover, as it was only in the direct condition that the force applied by the active finger would naturally cause a tactile sensation in the passive finger, the authors suggested that this attenuation was based on an internal prediction of the expected sensory consequences of the action. In the indirect condition the relationship between movement of the active finger and force in the passive finger was mediated by the response device in a novel manner that has previously been shown to impair sensorimotor prediction [12].

These results have been replicated and the predictive account corroborated in a range of studies that have investigated the sensory attenuation phenomenon using different experimental settings and manipulations [3, 4, 13]. Nevertheless, a detailed understanding of the computational basis of sensory attenuation has remained elusive. In particular, previous studies have debated whether attenuation is based on a subtractive mechanism, in which the sensation of an external force exceeds that of a self-generated force by a fixed amount irrespective of the force magnitude, or a divisive mechanism, in which the level of attenuation scales with the amplitude of the predicted force [5, 14]. This is significant because a subtractive mechanism could operate with only a coarse expectation that a self-generated force will be felt, whereas a divisive mechanism implies a fine prediction of the force amplitudes resulting from self-action.

While models of sensory attenuation have been presented in previous work at varying levels of computational detail [1, 15–17], these models have not addressed the perceptual and cognitive demands specific to force reproduction. Here we present the first detailed model of the computations that lead to exaggeration of target forces in the force matching task. We first validated the model framework by testing a prediction about trial-to-trial variability in matching force in new and previously published data sets. Our hypothesis was that the variability of the matching force would be greater in the direct condition compared to the indirect one, even when the mean matching force was equated. We then formally compared model variants with divisive and subtractive mechanisms, finding decisive evidence for a divisive attenuation process. Our findings expand our understanding of the attenuation process and provide further evidence against non-predictive gating [18–20] as an account of force matching responses.

## MATERIALS AND METHODS

### Model

Our model of the processes involved in the force matching task is set out in Fig. 1D. The black path is involved in both direct and indirect conditions while the blue path is unique to direct force generation. The filled circles represent the observable quantities: the target force exerted by the torque motor (*F*_*T*_) and the matching force exerted by the participant (*F*_*M*_). These are the only measured variables in the experiment. Perceptual noise corrupts the participant’s tactile perception of both the target force and the matching force (*s*_*T*_ and *s*_*M*_ respectively). Moreover, after experiencing the target force, the participant has to hold that sensation in memory (*m*_*T*_) for comparison with the matching force, and this process may introduce additional variability and bias (memory noise).

During the reproduction phase, the participant adjusts their output force *F*_*M*_ until the sensation of the force in the passive finger 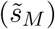 is equal to the remembered sensation of the target force *m*_*T*_, again with the possibility of some inherent variability (comparison noise). In the direct condition, an efference copy of the motor output from the active hand (*f*_copy_) provides input to a forward model that predicts the tactile sensation it will produce on the passive finger (*s*_pred_). This prediction is combined with the afferent sensory signal *s*_*M*_ to produce an attenuated sensation 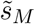. In the indirect condition, where the causal relationship between active finger movement and passive finger sensation is unnatural, we assume the sensation of the matching force is not attenuated, i.e., 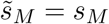.

This formal model identifies two important features of the force matching task. First, consistent differences observed between target and matching forces cannot be straightforwardly ascribed to the attenuation process, as they could also reflect biases in memory of the target force. Therefore, rather than compare matching forces to the target force, we can compare matching forces generated in direct and indirect conditions in response to the same target force. Such a comparison has identical memory and decision components and therefore allows us to isolate the attenuation process. Second, the sources of noise on the predictive pathway (blue) are expected to produce trial-by-trial variability in attenuation, which should translate into greater variability in the matching force when the attenuation process is active, i.e., in the direct condition as compared to the indirect condition. Based on these features, we analyzed data from the force matching task, focusing on differences between direct and indirect conditions in mean amplitude and trial-to-trial variability of the matching force.

### Data

In this study we combined analysis of new data and re-analysis of previously published data. Specifically, we included four data sets from experimental studies on force matching tasks. We extended a study by Kilteni and Ehrsson [21] by adding new data from 93 participants to the existing data set of 28 participants. The experimental protocol was identical for all 121 participants and is summarized below; see [21] for full methodological details. The other three data sets were from previously published studies on this topic [5, 22, 23].

Table 1 outlines sample characteristics and methodological details of each study. In all cases, target force levels were uniformly distributed across the stated range and tested with equal frequency in a pseudorandomized order of trials. On each trial, participants were given 3 seconds to reproduce the target force and the matching force was computed as the mean force exerted in the time interval between 2 and 2.5 seconds after the go signal.

**Table 1:**
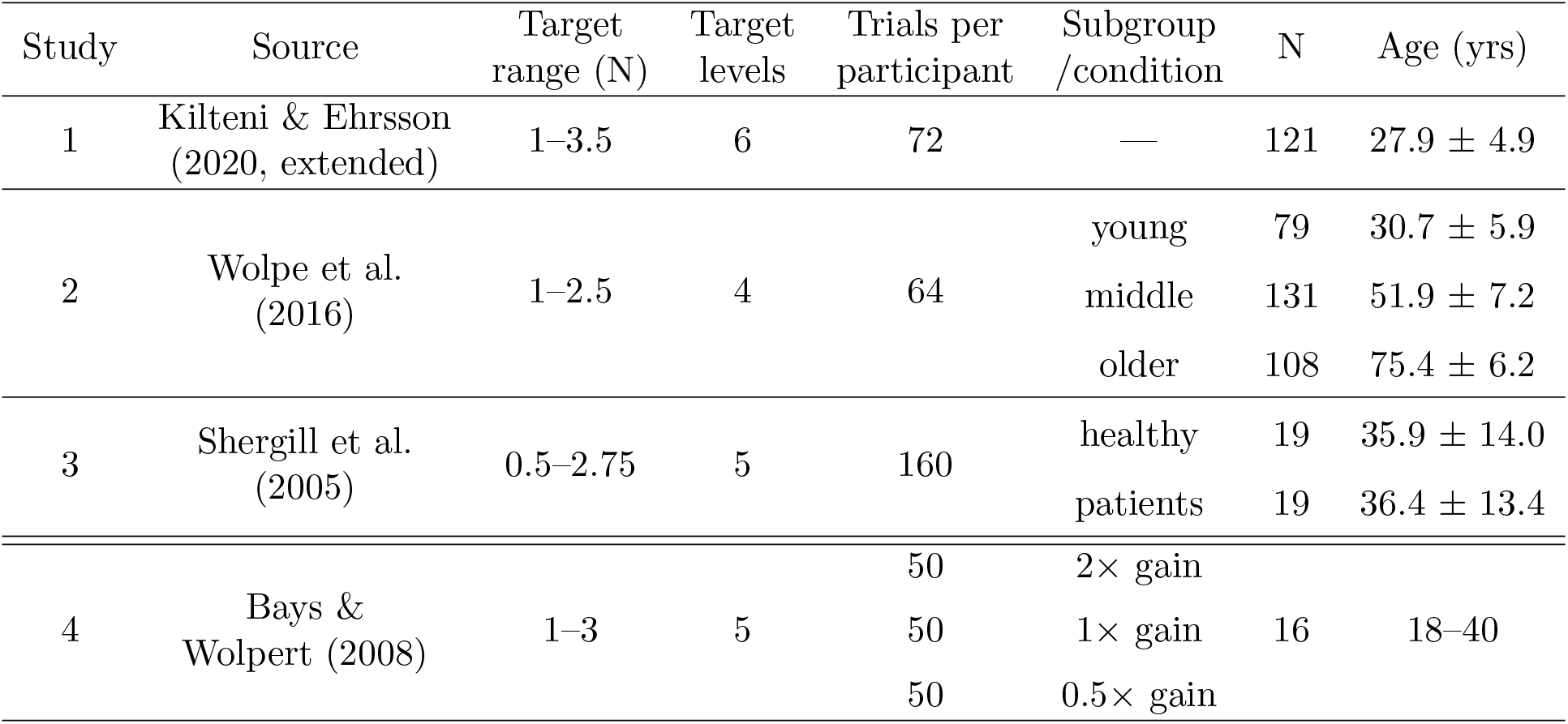
Summary of data sets

In Studies 1–3, trials were equally divided into blocks of direct and indirect force matching, with Studies 1 & 2 using a linear potentiometer (or “slider”) and Study 3 using a joystick to control force output in the indirect condition. In the direct condition, participants in Studies 2 & 3 pressed on the top surface of the lever to reproduce the target force, while in Studies 1 & 4 they pressed on a force sensor positioned above the lever while the torque motor continuously transmitted the recorded force to the passive finger, closely mimicking direct contact between the fingers. The force was always transmitted veridically in Study 1, whereas trials in Study 4 were equally divided into three blocks in which the gain between sensed and transmitted force was varied, so the force generated on the passive finger was 2*×* (double), 1*×* (veridical) or 0.5*×* (half) that of the participant’s active finger press. As a consequence, the force generated by, and tactile feedback received in the active finger varied across gain conditions while participants produced approximately the same matching force in the passive finger.

We excluded from the analyses four participants from Study 1 and two participants from Study 2 since they had missing data and two participants from Study 2 since the matching forces they produced did not correlate with the target forces, suggesting they likely did not properly understand the task.

Participants in Studies 2 & 3 were divided into subgroups that were analyzed separately. Participants from Study 2 were part of the Cambridge Centre for Ageing and Neuroscience (Cam-CAN) [24, 25] cohort of healthy adults spanning the adult age range, and were separated by age into young adult, middle-aged and older adult subgroups following the same criteria as the source study [23]. Study 3 comprised a patient subgroup with schizophrenia and an age-matched healthy control subgroup.

Studies 1–3, although using slightly different apparatus and including different populations, investigated the sensory attenuation phenomenon using the same force matching task, allowing their results to be directly compared. Replicating our analyses across these three independent data sets strengthens the results and demonstrates generalization of our findings. In contrast, Study 4 was analyzed and is presented separately from Studies 1–3, as it did not have an indirect force matching condition, instead having a within-participant manipulation of gain that made it specifically suited for evaluating the gating account of attenuation.

### Analysis

To assess qualitative predictions of the model framework for direct versus indirect force matching, we first computed, for each participant in each of Studies 1–3, the mean 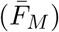 and standard deviation 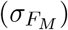 of their matching forces over all trials with each target force level (*F*_*T*_), separately for each condition. We then performed a linear mixed-effects model (*fitlme* in MATLAB) with subject as a random factor on these measures to generate lines of best fit and estimates of slopes and intercepts. Study 4 was omitted from this analysis as it had no indirect force matching condition.

When performing statistical inference on regression parameters, we used the Bayesian approach implemented in MATLAB (https://klabhub.github.io/bayesFactor/) with the default Jeffreys-Zellner-Siow prior on effect sizes [26]. The reported Bayes factors compare the predictive adequacy of two competing hypotheses (e.g., alternative and null) and quantify the change in belief that the data bring about for the hypotheses under consideration [27]. For example, BF_10_ = 10 indicates that the data are ten times more likely to occur under the alternative hypothesis (i.e., there is a difference) than under the null hypothesis (i.e., there is no difference). Evidence for the null hypothesis is indicated by BF_10_ *<* 1, in which case the strength of evidence is indicated by 1*/*BF_10_.

### Formal model specification

For quantitative fitting, we developed two implementations of the attenuation model illustrated in Fig. 1, incorporating either a subtractive or divisive mechanism.

### Subtractive model

According to the subtractive model, the tactile sensory attenuation is fixed, irrespective of the intensity of the stimulus. That is, given an afferent sensory signal *s*_*D*_, the participant perceives an attenuated sensation 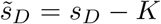, where *K* is the subtractive attenuation. We assume *K* is normally distributed with mean 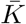 and standard deviation *σ*_*K*_. We further assume that, for a given target force, the matching force in the indirect condition, *F*_*I*_, is normally distributed with mean 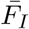 and SD 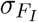, respectively. This matching force contains all the components of perception for the direct condition except for any attenuation. Consequently the matching force in the direct condition, *F*_*D*_, for the same target force will also be normally distributed with mean and SD,

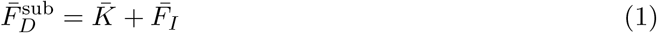

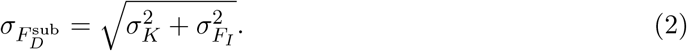

In contrast to the divisive model (below) the skewness of matching forces predicted by the subtractive model is always zero.

### Divisive model

The divisive model is based on the hypothesis that the attenuation is proportional to the stimulus intensity, i.e. 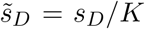. Again assuming *K* and *F*_*I*_ are normally distributed, the matching force in the direct condition is the product of two normally distributed variables. This distribution function does not in general have a closed-form expression, but can be approximated by a skew-normal distribution matched for mean, SD and skewness ([28]; see Supplementary Information for derivations),

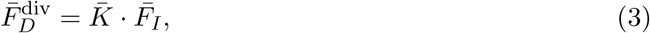

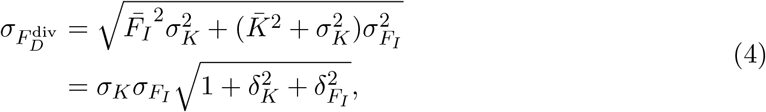

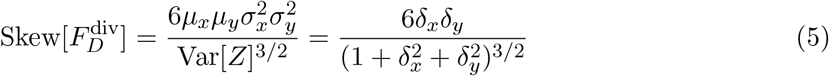

where we have used, 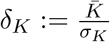 and 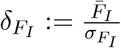.

### Hierarchical Bayesian modeling

We employed hierarchical Bayesian estimation to fit the models to data [29, 30]. In this modeling framework, a unique set of parameters is estimated for each individual participant, but all instances of each parameter are considered samples from a population-level distribution. Consequently, the estimation of each individual-level parameter is simultaneously informed by data from all the other participants – this occurs because all individual data contribute to the estimation of the population-level parameters, which, in turn, constrain all the individual-level parameters. Therefore, the hierarchical models offer a twofold advantage compared to traditional fitting approaches: these models share informative power across model hierarchy, which is particularly beneficial in scenarios with limited data (e.g., a low number of trials per participant, as is the case in some of the data sets we examined), and also for regularization of parameter estimates across individuals, enhancing the reliability of these estimates [31]. An important feature of the Bayesian approach is that it provides a posterior distribution over potential parameter values, rather than just point estimates of the best-fitting parameters as is the case with classical modeling approaches (e.g., ML). This full posterior distribution not only offers insights into the best point estimate (e.g., MAP) but also provides information about the uncertainty of parameter estimation (e.g., the width of the posterior distribution).

### Data fitting

We fit subtractive and divisive models (see Formal Model Specification) separately to each subgroup in each of Studies 1–4 (Table 1). We simultaneously estimated the posterior distributions of parameters by using the individual-trial data from all conditions within each experiment. Detailed model descriptions, along with specified prior distributions, can be found in the Supplementary Information. In brief, we fit a separate set of parameters for each input force level in the indirect force reproduction task. Using those measured matching forces, we estimated the attenuation parameters to fit the direct force reproductions.

To obtain the hierarchical fit, we used the Gibbs sampler with Markov Chain Monte Carlo algorithm implemented in JAGS [32], in combination with the *matjags* package [33] for MATLAB (The Mathworks Inc.). Posterior distributions for all parameters were obtained by running three parallel chains, with a thinning factor of two, and discarding the initial 1,000 samples for burn-in, resulting in a total of 10,000 valid samples per chain. After obtaining the samples for each dataset and model, we performed standard checks for convergence of chains (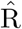; [34]) and autocorrelation within chains, and found the satisfactory performance of the sampler for both individual and population-level parameters. To compare how well the two competing models fit the data, we used DIC (Deviance Information Criterion) as a comparison metric. DIC is a generalization of Akaike’s Information Criterion (AIC), and it similarly depends on the goodness of fit and model complexity. The model with a smaller DIC is expected to be the one that would best predict and replicate the data.

We summarize the obtained posterior distributions numerically by reporting the median of posterior samples and the 90% highest density interval (HDI). The 90% HDI includes parameter values that have at least some minimal level of posterior credibility as opposed to values outside the HDI [35]. Specifically, the probability density is greater for any parameter value within the 90% HDI compared to values outside it, and the total probability of values within the 90% HDI is 90%. Consequently, the width of the HDI serves as a measure of the uncertainty associated with the parameter estimates, i.e., when the HDI is narrow, we can be relatively certain about the parameter estimates.

When analyzing differences in parameter values between different subgroups of participants, we first calculate group mean difference (MD) by finding the difference between the posterior distributions of two groups. We then report 95% HDI of the difference scores to quantify a range of values that has a 95% probability of containing the true difference in parameter estimates for two groups. Because parameter values with higher density are interpreted as more credible than parameter values with lower density, we set a decision rule to reject a mean difference of zero (i.e., no difference) when the 95% HDI does not contain zero. Intervals that include zero suggest zero as a credible difference value, indicating no difference between groups.

To assess the descriptive adequacy of the model, we generated posterior predictive distributions. The posterior predictive distribution shows what data to expect given the model at hand and the available knowledge, which includes the posterior distribution over parameters informed by the observed data [30]. The posterior predictive distribution is a distribution over data and therefore indicates the relative probability of different outcomes (i.e., output forces) after the model has seen the data. By comparing this predicted data to the data already observed, we can evaluate the model’s adequacy. We generated posterior predictive distributions of output forces in both direct and indirect reproduction conditions by using randomly sampled parameters from individual-level posterior distributions and the generative process described for each model. We conducted 10,000 simulations of output forces for each participant and condition. We then assess the resemblance between simulated and observed data through a combination of qualitative and quantitative methods. Qualitatively, we visually examine graphical displays of observed and simulated data.

Additionally, we quantitatively assess the fit by calculating the Bayesian *p*-value (not to be confused with the frequentist *p*-value). This value quantifies the probability that the obtained parameters will produce simulated data that deviates more from the predicted (average) data than the observed data does [36]. The Bayesian *p*-value of 0.5 indicates that observed and simulated data deviate equally around the model’s prediction. If the model’s predictions exhibit bias, meaning they are too extreme in either direction, the values will tend to deviate from 0.5, approaching zero or one. In such cases, one should consider rejecting the model and exploring alternative models. Given that the tested models are designed to overfit the data in the indirect reproduction conditions and hence expected to produce predictions very similar to the data, we focus on Bayesian *p*-values in the direct reproduction condition.

## RESULTS

We first examined whether the predictions of the model were qualitatively consistent with behavior in three existing datasets where participants performed force matching in both direct and indirect conditions. Fig. 2A illustrates the typical pattern of matching forces using data from Study 1 as an example (Kilteni & Ehrsson; see Fig. A2 and Fig. A3 for other datasets). The relationship between target force and mean matching force in each condition was close to linear, as shown by lines of best fit in Fig. 2A (linear regression model *R*^2^ = 0.94; for all subgroups of Studies 1–3, *R*^2^ *>* 0.91). In line with typical findings, directly generated matching forces (cyan symbols) consistently exceeded target forces (dashed line of equality) at every force level. As a function of target force, the matching force showed increases in both slope (*>* 1, BF_10_ = 1.92 *×* 10^18^) and intercept (*>* 0, BF_10_ = 3.35 *×* 10^9^) (cyan bars in Fig. 2C), a pattern that was also observed in Study 2 [23] for every age-based subgroup (young subgroup: slope BF_10_ = 9.15 *×* 10^6^, intercept BF_10_ = 1.58 *×* 10^5^; middle-age subgroup: slope BF_10_ = 1.22 *×* 10^14^, intercept BF_10_ = 1.21 *×* 10^11^; older group: slope BF_10_ = 2.64 *×* 10^4^, intercept BF_10_ = 3.74 *×* 10^9^). The healthy subgroup in Study 3 [22] showed a significant increase in intercept only (BF_10_ = 6.63 *×* 10^5^; slope BF_10_ = 0.24), while the patient subgroup (who displayed reduced attenuation overall according to the original study) showed an increased intercept (BF_10_ = 2.14 *×* 10^8^) paired with a decrease in slope (BF_10_ = 7.22).

**Figure 2:**
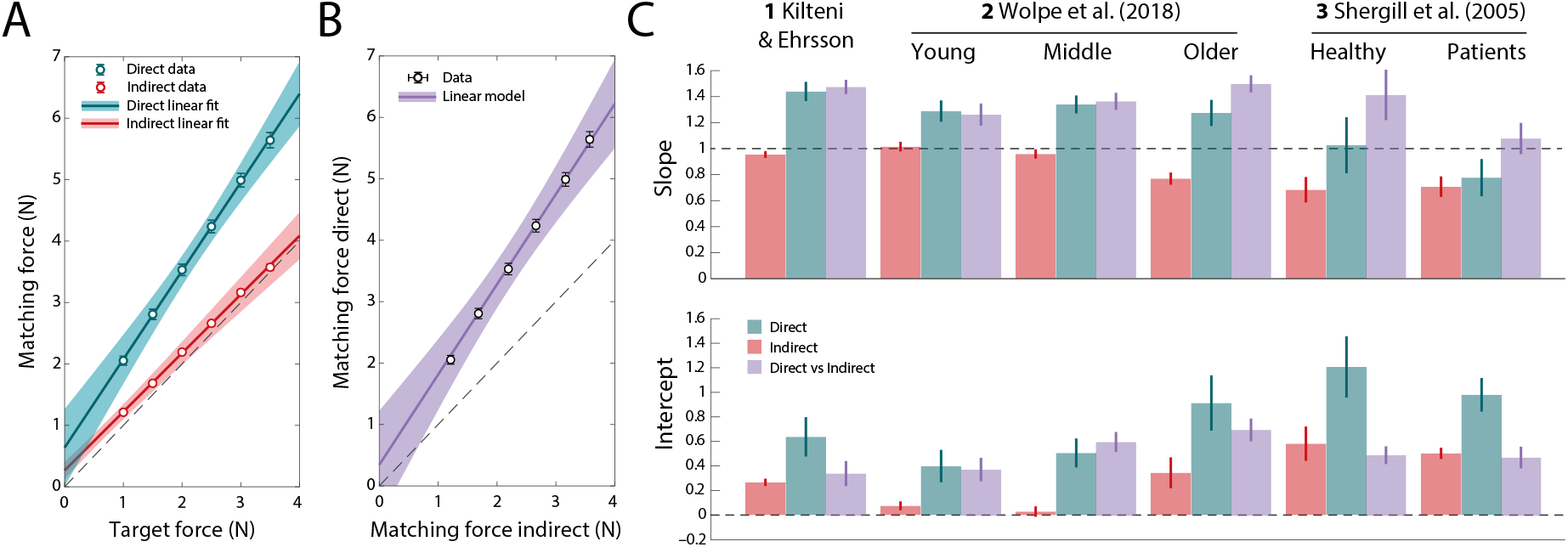
Matching forces in direct and indirect force reproduction. (A) Mean matching force as a function of target force in Study 1, for direct (cyan) and indirect (red) conditions. Data are shown as symbols and linear fits as lines. (B) Matching force in the direct condition of Study 1 plotted against matching force in the indirect condition, with linear fit. Symbols indicate mean matching forces for each target force level. (C) Slope (top) and intercept (bottom) parameters for each data set and subgroup, for the regression of direct (cyan) and indirect (red) matching force against target force (as in A) and direct matching force against indirect matching force (as in C; purple). Error bars in A & B indicate *±*SE. Shaded areas in A & B and error bars in C indicate 95% CI. In A & C (cyan and red), slope greater than one indicates exaggeration of the matching force proportional to the target force magnitude, while intercept greater than zero indicates exaggeration of the matching force irrespective of the target force magnitude. In B & C (purple), slope greater than one means that the matching force in the direct condition was proportionally higher than the matching force in the indirect condition, while intercept greater than zero indicates a systematic offset between direct and indirect matching forces.

Consistent with classical findings, reproduction of target forces was substantially more accurate when the matching force was generated indirectly (red symbols in Fig. 2A). Nonetheless, in Study 1, linear regression coefficients indicated deviations from equality in both slope (*<* 1, BF_10_ = 20.55) and intercept (*>* 0, BF_10_ = 2 *×* 10^31^). This pattern was replicated in the other studies and subgroups (red bars in Fig. 2C), with the exception of the young adult subgroup (strong evidence for an increase in intercept, BF_10_ = 203; moderate evidence against a change in slope, BF_10_ = 0.17) and the middle-aged subgroup (weak evidence for a reduction in slope, BF_10_ = 1.13; moderate evidence against a change in intercept, BF_10_ = 0.23) of Study 2. These results are consistent with predictions of our model of the task (Fig. 1D), in which accumulated bias and variability in memory for the sensation of the target force, possibly supplemented by noise in the comparison process itself, cause matching forces to deviate from target forces even in the absence of predictive attenuation. More specifically, the particular pattern observed across studies for the indirect condition, of a decrease in slope paired with an increased intercept, is consistent with the classical finding of contraction bias in memory [37, 38], whereby memory for the current target force is influenced by the history of target forces on preceding trials, leading to a bias towards the mean of the presented forces. Notably, in Study 2 we observed a progressive flattening of indirect matching slopes with age, consistent with the well-established age-related decline in short-term memory performance [39, 40].

### Isolating effects of predictive attenuation

The results above are typical of a wide range of previous force matching studies, in which healthy participants were found to be close to accurate in indirect force reproduction, but substantially exceeded target forces in direct reproduction. However, when analyzing slopes and intercepts there is considerable variation across groups and studies, making it difficult to discern a consistent pattern. We hypothesized that this variation was at least partly due to the memory and comparison components identified in our model of the task, rather than the attenuation of perceived force in the direct condition. Because these non-predictive components are identical in direct and indirect conditions, the effects of predictive attenuation should be better isolated by comparing matching forces in the two conditions for the same target force. Fig. 2B plots data from Study 1 in this way (see Fig. A2 and Fig. A3 for other datasets). This relationship was again close to linear (*R*^2^ = 0.97; for all subgroups of Studies 1–3, *R*^2^ *>* 0.96), with slopes and intercepts plotted as purple bars in Fig. 2C. Relative to results from the individual conditions (cyan and red bars), comparing direct to indirect matching forces reduced variability across subgroups, revealing consistent evidence for a strong deviation in slope in every subgroup (excepting patients in Study 3) and a relatively reduced (though still non-zero) intercept.

This approach also clarified results for the patients in Study 3 (rightmost bars in Fig. 2C), indicating that the greater accuracy of force matching in patients with schizophrenia reflected a strong reduction in slope compared to age-matched controls, approaching veridicality (slope of 1), but negligible change in intercept. In Study 2, the increasing exaggeration of target forces with age was observed in both slope and intercept, once memory effects were accounted for.

### Evidence for excess variability in direct force reproduction

Our model of force matching predicts excess variability in the direct compared to indirect condition (Fig. 1D), arising from variability in the predictive processes involved in sensory attenuation (i.e., efference copy noise and prediction noise in Fig. 1D). Fig. 3A plots within-participant across-trial variability in matching force as a function of target force level for Study 1. Matching force variability was substantially greater in the direct condition (cyan) than the indirect condition (red) at each target force level compared to the indirect condition. This finding was consistent across all data sets and subgroups as measured by differences in regression parameters between the two conditions (Fig. 3B; all BF_10_ *>* 5.6).

**Figure 3:**
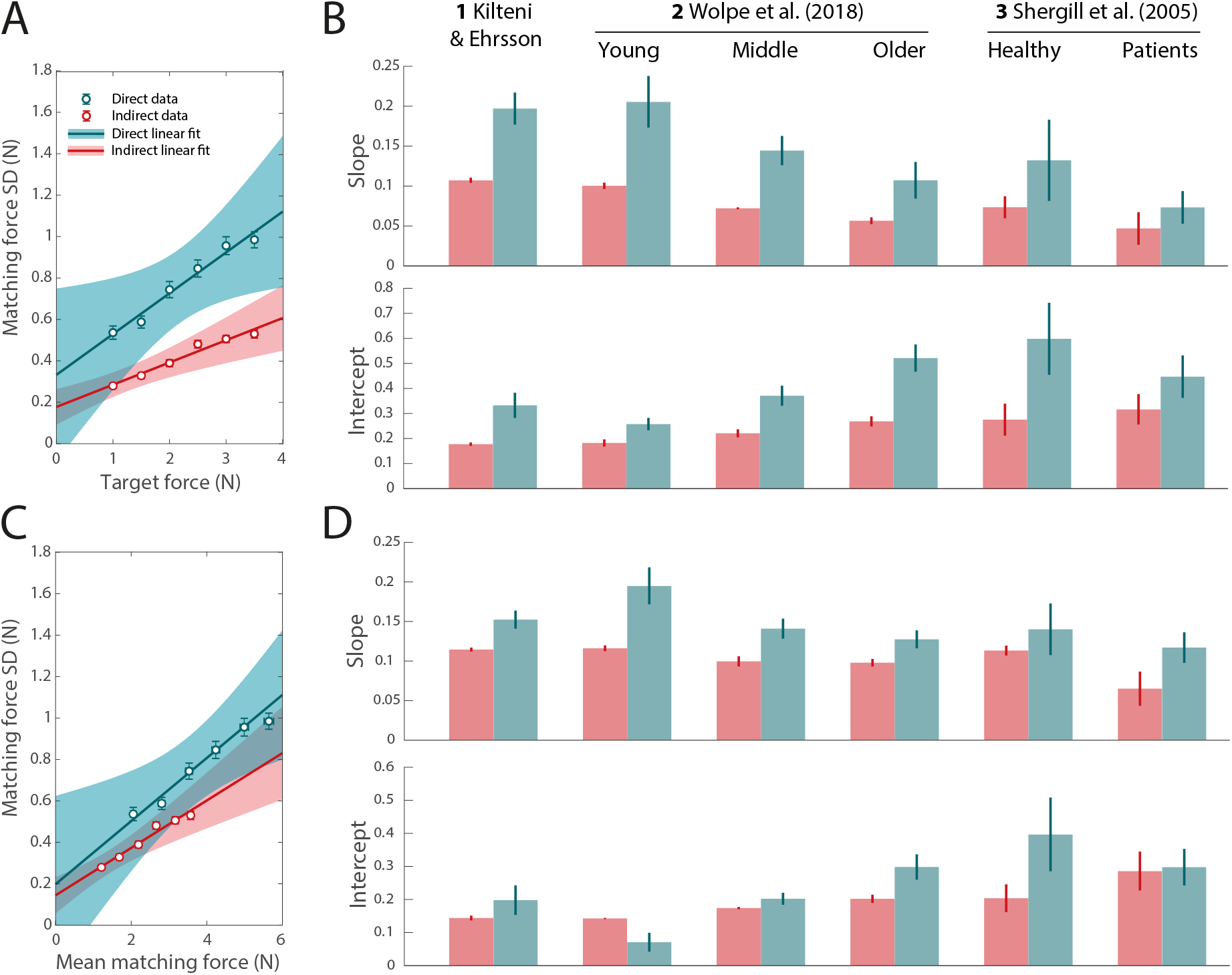
Trial-to-trial variability in matching force. (A) Average of within-subject standard deviation of matching force as a function of target force in Study 1, for direct (cyan) and indirect (red) conditions, with linear fits. (B) Slope (top) and intercept (bottom) parameters for each data set and subgroup, for the regression of direct (cyan) and indirect (red) matching force SD against target force (as in A). (C) Average of within-subject standard deviation of matching force as a function of mean matching force in Study 1, with linear fits. Each symbol corresponds to a single target force level. (D) Parameters of the linear regression of matching force SD against mean matching force (as in C) for each data set and subgroup. In A & B, slope greater than zero indicates that matching force variability increases with increasing target force, while intercept greater than zero indicates the presence of trial-to-trial variability irrespective of the target force magnitude. In C & D, slope greater than zero indicates that matching force variability increases with increasing mean matching force, while intercept greater than zero indicates the presence of trial-to-trial variability that does not depend on the mean matching force magnitude. Error bars in A & C indicate *±*SE. Shaded areas and error bars in B & D indicate 95% CI.

Although matching force variability was consistently larger in direct than indirect condition for the same target force, the mean matching force were also higher in the direct condition. Variability of force output has been found to scale with the mean amplitude [41, 42], so to examine whether this was a viable explanation for the increased variability we next regressed matching force SD in the two conditions against mean matching force (Fig. 3C shows results from Study 1; Fig. 3D plots regression coefficients for each dataset and subgroup). As predicted, variability increased with mean force amplitude in both direct and indirect condition (positive slope in all subgroups; all BF_10_ *>* 1644). However, matching force variability was consistently greater in direct than indirect condition for the same mean force amplitude (cyan symbols above red symbols in Fig. 3C), an effect primarily driven by increased slope in the regression model for every dataset and subgroup (cyan bars vs. red bars in Fig. 3D, top; healthy subgroup BF_10_ = 1.1, all other BF_10_ *>* 591).

### Subtractive versus divisive models of attenuation

Having confirmed qualitative predictions of the model framework, we next computed formal model fits for each of the data sets and subgroups. This required us to specify how the attenuation process quantitatively changes the sensation of force: we considered two variant models, one in which the attenuation was subtractive and one in which it was divisive. In the subtractive model, the attenuation process subtracts a fixed amount (the attenuation factor) from a perceived force only when it is directly self-generated: as a result the matching force in the direct condition exceeds that in the indirect condition by a fixed amount. In the divisive model, a fixed fraction of a perceived force is instead removed when self-generated, equivalent to dividing it by the attenuation factor: as a result the matching force in the direct condition equals the matching force in the indirect condition times the attenuation factor. In both variant models, the attenuation factor was fixed with respect to different levels of target and matching force but incorporated trial-to-trial variability. Specifically the attenuation factor was modelled as a normally distributed random variable with mean and SD as free parameters.

### The divisive model provides a close fit to behavioural data

Formal model comparison demonstrated that the divisive model provided a superior fit to behavioural responses on the force matching task compared to the subtractive model in all subgroups of Studies 1–3 (Study 1, ΔDIC = 1039.4; Study 2, ΔDIC ≥ 201 for all age-groups; Study 3, healthy participants: ΔDIC = 169.1, patients: ΔDIC = 2.5). Below, we present further results obtained from fitting the divisive model. Results for the subtractive model are available in the Supplementary Information (Table A1; Fig. A6 & Fig. A7).

The fit of the divisive model to data is exemplified by the posterior predictive distributions for Study 1 shown as shaded areas in Fig. 4. These model predictions accurately reproduced the empirical distributions (grey histograms) of matching forces in direct (cyan) and indirect (red) conditions at each force level. The effect of attenuation in the direct condition is observed as increases compared to the indirect condition in mean, variance and skewness of the matching force distributions. Skewed distributions (note long tails in direction of larger matching forces in direct condition; Fig. 4) are a prediction of the divisive model, arising from multiplication of two unskewed random variables (attenuation factor and indirect matching force), and contribute to its superior fit over the subtractive model.

**Figure 4:**
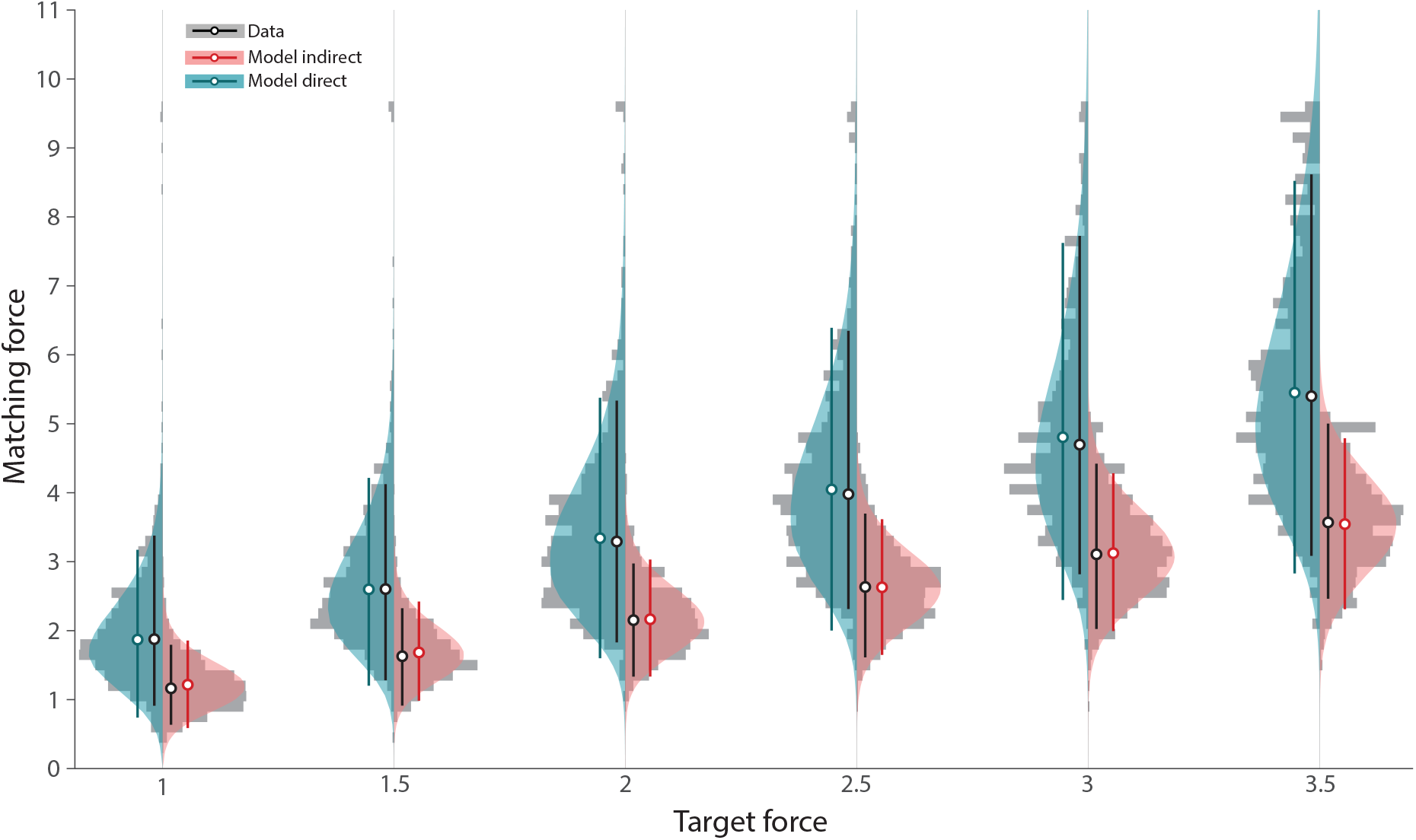
Posterior predictive densities for Study 1. Grey histograms show matching forces pooled across all participants. Coloured distributions show simulated matching forces based on posterior predictive density. The circles with error bars show the median and 90% HDI for data (black) and simulated trials for the same condition (cyan: direct, red: indirect).

To quantify the quality of fit, we calculated Bayesian *p*-values from the posterior predictive distributions in the direct condition. We observed Bayesian *p*-values that were consistently around the ideal value of 0.5 (0.47 ≤ all *p* ≤ 0.51), indicating a close match between simulated and observed data. The divisive model provided a similarly close match to the data from Studies 2 & 3 (Fig. A4 and Fig. A5, respectively). Again, the strong resemblance between the simulated and observed data was numerically verified by calculating the Bayesian p-values for each dataset and target force level, consistently finding values around 0.5 both in Study 2 subgroups (young 0.46 ≤ all *p* ≤ 0.52; middle 0.47 ≤ all *p* ≤ 0.53; older 0.47 ≤ all *p* ≤ 0.52), and Study 3 subgroups (healthy controls: 0.45 ≤ all *p* ≤ 0.52) and patients: 0.44 ≤ all *p* ≤ 0.54).

### Mean and variability of the attenuation factor

Figure 5A displays the observed posterior distributions for the population-level mean attenuation factors from the divisive model. We summarized these posterior distributions by calculating the median values. In Study 1, the estimated (posterior median) mean attenuation factor was 1.61 (90% Highest Density Interval, HDI =[1.56, 1.66], describes the range of values within which the parameter of interest lies with 90% probability). This value indicates that, on average at the population level, a force generated indirectly via the response device was perceived as 61% more intense than the same force when directly self-generated; or equivalently, that perception of the directly self-generated force was attenuated by 38% (1 − 1*/*1.61 = 0.38). Across the different subgroups of Studies 1–3, the posterior distributions of the mean attenuation factors exhibited clear heterogeneity. Importantly, none of the reported HDIs include the value of 1, indicating that in all examined datasets the perception of force in direct self-generation was attenuated in comparison to the indirect condition.

**Figure 5:**
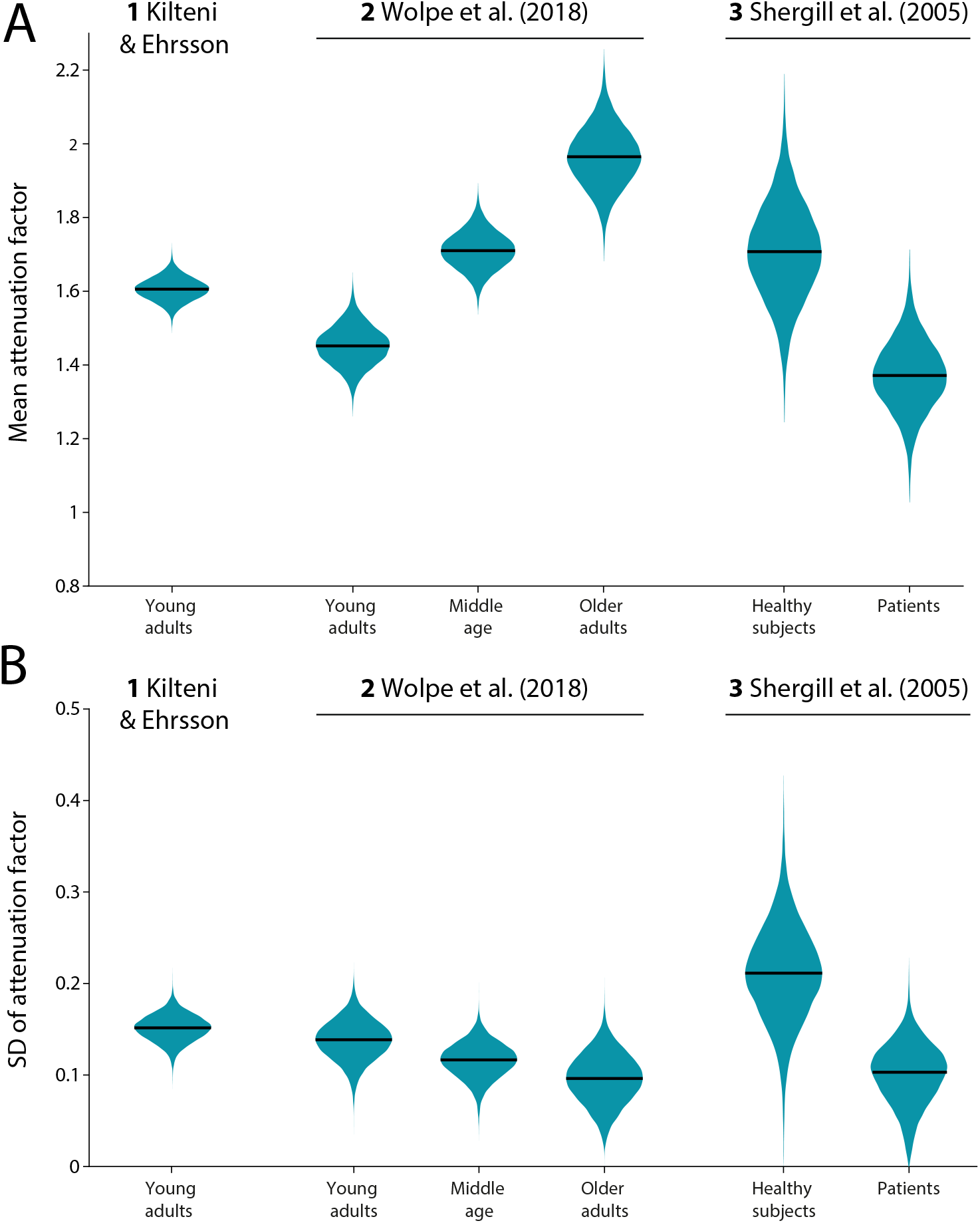
Posterior densities for (A) the mean attenuation factor and (B) the SD of attenuation factor, based on population-level central tendency parameters of the divisive attenuation model fit separately to each dataset and subgroup. Black lines indicate the posterior median.

The posterior distributions of standard deviation of the attenuation factor in the divisive model are shown in Fig. 5B. In contrast to the mean attenuation factor, estimates of this parameter exhibited much greater uniformity between datasets. Specifically, in Study 1, the median parameter was found to be 0.152 (90% HDI [0.128, 0.174]), indicating that the perceived intensity difference between conditions varied from trial-to-trial about its mean of 61% with an SD of 15%. This is approximately equivalent to an attenuation of the directly self-generated force by 38% *±* 6%. Similar values were found for the different subgroups of Studies 2 & 3, described in detail below.

### Attenuation scales with age

Study 2 compared the mean exaggeration of the target force in the direct condition across the three groups based on age difference, providing an opportunity to examine how sensory attenuation varies with age. Estimated mean attenuation factors were 1.45 (90% HDI [1.365, 1.536]) in the sample of young adults, 1.71 (90% HDI [1.63, 1.793]) in the sample of middle-aged participants, and 1.97 (90% HDI [1.834, 2.096]) in the sample of older adults (Fig. 5A). This suggests an increase in mean attenuation factor with age, consistent with conclusions of the source study. To corroborate this observation statistically, we calculated the mean differences (MD) between posterior distributions from these groups. We found that young adults had a lower mean attenuation factor compared to both middle-aged adults (MD = –0.259, 95% HDI [–0.402, –0.12]) and older adults (MD = –0.514, 95% HDI [–0.698, –0.322]), while middle-aged participants had a lower estimate compared to older adults (MD = –0.255, 95% HDI [–0.438, –0.067]).

In contrast, the standard deviation of the attenuation factor exhibited greater similarity among the three subgroups, with a trend to decreasing SD with age (Fig. 5B). To quantify this, we again calculated MDs between posteriors in different groups and, in all cases, found that the 95% HDI encompassed zero, indicating no evidence for a difference in estimated parameters. Specifically, this was the case when comparing young and middle-aged adults (MD = 0.022, 95% HDI [–0.033, 0.079]), young and older adults (MD = 0.042, 95% HDI [–0.025, 0.11]), and middle-aged and older adults (MD = 0.02, 95% HDI [–0.045, 0.082]).

### Patients with schizophrenia have reduced sensory attenuation

The data in Study 3 comprised groups of schizophrenic patients and neurotypical controls. Estimates of the mean attenuation factor were 1.708 (90% HDI [1.498, 1.928]) for controls and 1.372 (90% HDI [1.213, 1.526]) for patients (Fig. 5A). This suggests that, on average, healthy controls exhibited stronger sensory attenuation. To confirm this, we calculated the mean difference between the posterior distributions, revealing that the attenuation factor was indeed statistically larger in the group of healthy participants (MD = 0.3361, 95% HDI [0.007, 0.649]).

Comparing the estimated standard deviation of the attenuation factor between the two groups (Fig. 5B), we found that a trend for smaller SD in patients did not reach conventional significance, as indicated by a 95% HDI of MD containing zero (MD = 0.111, 95% HDI [– 0.008, 0.234]). Similarly to the comparison between samples in Study 2, a clear difference in the estimated attenuation factors was not accompanied by a significant difference in their variability. Note however that the posterior estimates from this study exhibited relatively higher uncertainty in parameter estimates (i.e., broader posterior) compared to other subgroups, most likely due to the smaller number of participants in each subgroup.

### Testing a non-predictive gating account of attenuation

A key component of the proposed model is a predictive mechanism (the forward model) that estimates sensory consequences of motor commands, enabling self-generated sensory inputs to be attenuated at an early stage of processing. Alternatives to this account propose that sensitivity to the matching force in the passive finger is decreased via a non-predictive gating mechanism triggered by muscle activation or tactile feedback in the active finger [18–20]. Such non-predictive gating has been clearly demonstrated in active or moving body parts [43–48], but not in passive ones. Because the right index finger is active in indirect as well as direct conditions (moving the slider or joystick), this account additionally requires that the gating is weaker in the indirect condition, perhaps because less force is required to adjust the response device.

To investigate the role of motor activity and feedback in the active finger in force matching, we applied our model to the data from Study 4, that manipulated the transmission of force from the active to the passive finger [5]. Fig. 6A illustrates forces produced by the active finger and forces transmitted to the passive finger across the target force levels in the three gain conditions.

**Figure 6:**
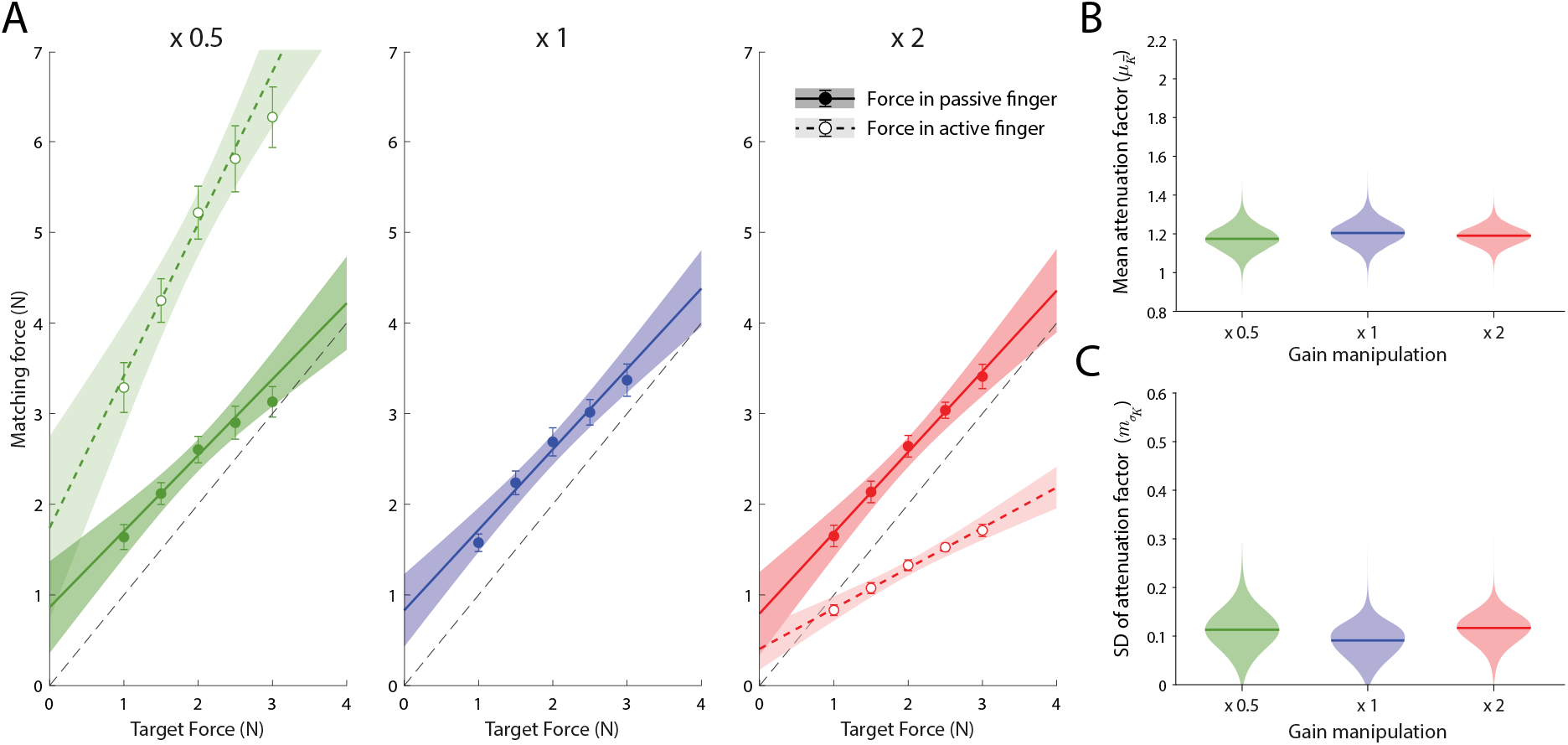
Evidence against a gating account. (A) Forces generated by active finger (empty symbols and dashed line) and forces applied to passive finger (filled symbols and solid line) during direct matching in the three different gain conditions of Study 4 (passive and active forces are identical for gain = 1, middle panel). Error bars indicate *±*SE; shaded areas indicate 95% CI. (B & C) Posterior densities for (B) the mean attenuation factor and (C) the SD of attenuation factor, based on the divisive attenuation model fit separately to each gain condition. Horizontal lines indicate the posterior median.

For the hierarchical fitting, as our interest was in comparing the three different conditions of direct reproduction, we fixed parameters corresponding to indirect force generation for each participant and condition to the same set of population-level parameters estimated from Study 1 (which had matching target force levels), and performed fitting on direct condition data separately for each gain condition. The observed population-level posterior distributions are shown in Fig. 6B. While the gating hypothesis would predict less attenuation as the factor multiplying the input force increases, we obtained highly consistent estimates of the mean attenuation factor across the three conditions. Specifically, the estimated (posterior median) mean attenuation factors were 1.175 (90% HDI [1.066, 1.281]) for the gain factor of 0.5, 1.204 (90% HDI [1.099, 1.312]) for the gain factor of 1, and 1.191 (90% HDI [1.108, 1.269]) for the gain factor of 2. We then calculated the mean differences between posterior distributions of these groups and, in all cases, found that the 95% HDI encompassed zero, indicating no differences in estimated parameters between these groups (MD_0.5−1_ = –0.03, 95% HDI [–0.211, 0.156]; MD_0.5−2_ = –0.016, 95% HDI [–0.176, 0.147]; MD_1−2_ = 0.014, 95% HDI [–0.148, 0.176]).

A similar conclusion can be drawn when comparing the posterior distributions of the standard deviation of the attenuation factor (Fig. 6C). The obtained posterior estimates were 0.113 (90% HDI [0.044, 0.18]) for the gain factor of 0.5, 0.091 (90% HDI [0.031, 0.138]) for the gain factor of 1, and 0.117 (90% HDI [0.065, 0.163]) for the gain factor of 2. Calculating MDs between these posterior distributions, here we also found that the 95% HDI in all cases encompassed zero, indicating no differences in estimated parameters between these groups (MD_0.5−1_ = 0.024, 95% HDI [–0.079, 0.129]); MD_0.5−2_ = –0.001, 95% HDI [–0.103, 0.099]); MD_1−2_ = –0.026, 95% HDI [–0.112, 0.064])).

## DISCUSSION

Here, we proposed a formal computational model of the processes leading to exaggeration of target forces in the force matching task [2]. According to this model, matching forces recorded in both direct and indirect reproduction conditions are subject to identical sources of perceptual, memory, and comparison noise, differing only in the presence or absence of predictive attenuation. On this basis, we isolated a pure measure of attenuation strength by comparing the matching forces between the two conditions. This analysis confirmed and extended previous conclusions that attenuation is enhanced in older compared to younger individuals [23] and weaker in patients with schizophrenia compared to age-matched control participants [22].

Previous studies suggested that a network including cerebellum, secondary somatosensory cortex (SII) and premotor cortex is involved in sensorimotor prediction and sensory attenuation (e.g. [21, 49, 50]). Reduced connectivity in cerebellar-midbrain and cerebellar-thalamic networks has been proposed as a plausible functional basis for the role of cerebellum in psychotic disorders like schizophrenia [51, 52]. The reduction in sensory attenuation previously observed in patients with schizophrenia [22], and further delineated by our analysis, could be related to impaired cerebellar function weakening the ability to predict movement consequences or use the prediction to attenuate matched sensory input. On the other hand, the increased attenuation found in older participants suggests that cerebellar impairment cannot provide a straightforward account of all modulations of sensory attenuation.

Removing the confounding effect of memory bias proved particularly important for older individuals: in indirect force matching we observed a bias towards the average target force that grew stronger with age. This is consistent with a classical contraction bias [37, 38] in which memory of the target force on the current trial is drawn towards the average of target forces on previous trials. The increase in bias with age may reflect a corresponding decline in working memory fidelity.

Our model made a specific prediction of excess trial-to-trial variability in the direct compared to the indirect condition, attributed to variation in attenuation level due to noise in motor prediction. We experimentally confirmed this prediction in every dataset, observing higher variability in the direct condition compared to the indirect condition, even when accounting for the different magnitudes of the mean matching forces. One possible counterargument is that, rather than prediction noise, the higher variability in the direct condition could stem from higher motor noise, since participants use their right index finger to press in the direct condition but use it to manipulate a response device in the indirect one. However, the force matching studies were designed to minimize contributions of motor noise by averaging the force applied to the passive finger on each trial over a period of 500 ms. Moreover, if motor noise was responsible for the excess variability, the higher motor variability typically observed in older adults [53] would then be expected to produce greater excess variability, whereas we observed comparable variability across all age groups. Similarly, the trend towards smaller excess variability observed in the patients with schizophrenia would imply that patients have lower motor variability compared to controls, contradicting previous findings [54].

We proceeded to test two alternative accounts of how attenuation affects predicted force sensations. According to a subtractive account of attenuation, a self-generated force is perceived as weaker than an externally generated force by a fixed amount that does not scale with the force amplitude. Supporting this notion, Walsh, Taylor, and Gandevia [14] and Bays and Wolpert [5] found that attenuation in the direct condition mainly consisted of a fixed “offset” from the target force. However, neither of these studies compared the direct condition to an indirect condition, raising the possibility that their findings might reflect biases unrelated to prediction, e.g., the above-mentioned contraction bias. According to the alternative divisive account, the difference in perceived force scales with magnitude of the force, with lighter forces being subject to less attenuation compared to stronger forces. In other words, these two alternative hypotheses would have different effects on the perception of the weakest tactile stimuli. Formal model comparison decisively supported this divisive account of attenuation, based on the differing predictions for the distributions of matching forces. The divisive account of attenuation straightforwardly explains why even the lightest self-generated touch remains perceptible, despite being attenuated relative to an equivalent external force.

Because the direct and indirect conditions differ in the amplitude of force applied by the active finger, it has been argued [18–20] that overestimation of forces in the direct condition could arise from a non-predictive mechanism based on this activity, similar to the sensory gating that suppresses tactile input in a moving effector regardless of its predictability [43], but now spreading to affect the passive finger of the other hand. Applying our modelling methods to previous data from a gain manipulation experiment [5] provided evidence against the gating hypothesis. Despite a four-fold variation across gain conditions in the force output in the active finger, we observed attenuation of similar magnitude and variability across all conditions. This result appears incompatible with non-predictive gating due to activity in the hand applying the force, which varied widely across conditions, but consistent with a mechanism that attenuates the sensation based on a prediction of the force received on the passive finger, which remained approximately constant across conditions. These results also appear to rule out any non-predictive mechanism based on tactile feedback in the active finger, which similarly varied four-fold in amplitude across gain conditions.

These findings align with studies that found attenuation and gating are governed by separate mechanisms [13, 55, 56], and a range of previous observations supporting a predictive basis for attenuation of self-generated touch. These include the finding that attenuation is present when fingers are expected to make contact, even if they fail to do so [4] (recently replicated in [6]), whereas attenuation is not observed when the same movement is made without an expectation of contact, or when equivalent forces are presented passively to both fingers [3]. The constant attenuation across different gain conditions implies that the predictive mechanism could update to reflect the new gain relationship, consistent with previous evidence for adaptability of the attenuation process to delays in force transmission [57, 58].

In a previous study based on the “free-energy” model of [59], Brown et al. [15] proposed that sensory attenuation is a necessary component of action. According to this account, voluntary movements are the result of an internal prediction that movement is occurring: the disparity between the proprioceptive feedback expected on this basis and the actual feedback is resolved by the muscular system generating the predicted movement. These authors argued that strong evidence from the other senses that movement is not occurring will prevent movement initiation, and proposed to resolve this by increasing internal uncertainty about these sensory signals around the time of the predicted movement. In order to account for the attenuation of self-generated forces observed in the force matching task, the authors further proposed that the perceived magnitude of a force is fixed to the 90% lower confidence bound on the internal estimate of magnitude, i.e., that forces that have higher uncertainty associated with them are perceived as weaker.

The free-energy account has some common elements with our proposed model of attenuation, in particular that the mechanism in both cases is based on predicting the sensory consequences of self-action. However there are also significant differences. In the present model, the goal of predicting sensory consequences is to down-weight them relative to sensations with an external cause; excess variability is an inevitable but incidental consequence of noise in the predictive process. In the free-energy model the goal is to increase internal uncertainty about whether a force is being self-generated, because this is necessary to permit a force to be self-generated: attenuation is an incidental by-product of this increase in uncertainty, solely due to their assumption that more uncertain forces are perceived as weaker. Uncertainty could be increased by injecting noise into sensory inputs, which would predict excess variability in estimating self-generated forces, however this does not appear to be a necessary component of the free-energy model, as uncertainty could instead be added to the prior or directly to the posterior without increasing variability. Predictions for variability in force estimation were not discussed in the previous paper.

Importantly, [15] modelled attenuation of self-generated forces in the context of force matching, but did not model the sequence of events in a force matching trial (in fact in [15]’s simulations, external forces were generated to match the lower confidence bound on a preceding self-generated force) or consider how variability in perception, memory or sensory predictions would affect reproduction of an external force. Attenuation in the free-energy model is due to a seemingly *ad hoc* assumption relating perception of sensation to uncertainty, which conflicts with the wider literature on sensory perception, including Bayesian models of perception [60]. In contrast, the principle that self-generated sensations are actively cancelled, as in the present model, is supported by neurophysiological studies in multiple model systems [7–10].

More recently, the free-energy model inspired two works where a robotic arm was controlled by a hierarchical recurrent neural network (RNN) designed to follow the free-energy minimization principle [16, 17]. The RNN was trained to make movements with feedback consistent with self-generated and externally generated contexts. After training, the precision of the sensory prior was reduced in the self-produced context and increased for externally generated sensations, similarly to [15]. However, these studies did not address the perceived intensity of force sensation and did not model the force matching task.

A key assumption of our model of the force matching task is that the attenuation mechanism is inactive in the indirect condition. This might seem counter-intuitive, given that the force in the passive finger is “self-generated”, in the sense that it is controlled by the participant moving the joystick or slider and, at least in principle, predictable based on movement of the active hand. In common with previous accounts [2, 5, 21, 22], we suggest attenuation is absent because the causal relationship between action and sensory consequence in this condition is novel and lacks an ecological mechanism. This interpretation is consistent with previous findings that physically separating the finger applying a force from the finger receiving it reduces the level of attenuation (although does not eliminate it [5]), and that forces that are predictable based on active finger movement but not associated with a contact event do not show attenuation [3, 6, 20]. Previous evidence that the predictive mechanism can adapt to changes in the causal relationship [57, 58] suggests that attenuation might develop in the novel context with sufficient exposure, but this may take considerably longer than a typical experimental session. Note that sensory prediction is not necessary for the task of force reproduction using a joystick or slider (or indeed a finger press), as this can be achieved with a simple feedback control loop.

A possible alternative account for the absence of attenuation in the indirect condition is based on the fact that selective attention is divided between two spatial locations (the passive finger is located under the force lever while the active finger is at the location of the response device), whereas in the direct condition the fingers are approximately co-located. Although focused attention is not generally associated with a reduction in perceived intensity, it might be argued that the forces in active and passive fingers are more difficult to disambiguate when they are co-located, and that this confusion in some way leads to a lessening of the perceived force. However, this account would be inconsistent with the substantial body of evidence for predictive attenuation in “tapping” tasks, where the task-relevant events all occur in the same physical location, and *test* taps are compared with externally applied *reference* taps of varying amplitude to determine the point of subjective equality [3, 4, 55, 57]. For example, in [3], when a self-generated force was produced by an active finger tapping on a passive finger, the perceived force in the passive finger was attenuated, but when equivalent forces were externally applied simultaneously to both fingers, no attenuation was observed, despite the spatial locations and postures of the fingers being identical.

Taken as a whole, the observations above imply that predictability of a sensory input is necessary but not sufficient for it to be attenuated. Predictive attenuation may have evolved to enhance sensitivity to sensations with an external origin, perhaps because they are more likely to indicate a threat or otherwise require a behavioural response. For this purpose, down-weighting inputs based purely on their predictability would be inadequate, as many sensory events with external causes can be predicted (e.g., the sound of a door shutting can be predicted if you observe someone push it). Moreover, attenuating predictable signals could impair our ability to learn further causal relationships involving them. If attenuation requires a determination of self (versus external) cause in addition to predictability, it may be related to a sense of agency [61]. Future studies could examine whether the adaptation of attenuation to changed causal relationships (e.g., delays) is coupled to increases in the perception of self-agency in the interaction.

## Acknowledgements

This research was supported by the European Research Council (ERC), starting grant TICK-LISHUMAN (101039152) to KK, and the Wellcome Trust, grant 106926 to PMB. NV thanks Professor Nicola Smania (University of Verona, Italy) for helpful and insightful discussions. We thank all authors of source studies and the Cambridge Centre for Ageing and Neuroscience (CamCAN; https://cam-can.mrc-cbu.cam.ac.uk) for sharing data. A preprint is available at https://doi.org/10.1101/2024.06.20.599826.

## Current addresses

NV is currently with the University of Verona, Department of Engineering for Innovation Medicine, Verona, 37134, Italy. IT is at the University of Zagreb, Faculty of Humanities and Social Sciences, Department of Psychology, Zagreb, Croatia.

## Author contributions

PMB contributed to conceptualization, methodology, project administration, visualization, and writing – original draft and revisions. NV contributed to methodology, data analyses, and writing – original draft and revisions. IT contributed to data analyses, visualization, and writing – original draft and revisions. ZG contributed to methodology and data analyses. DMW contributed to conceptualization, methodology and writing - revisions. KK contributed to methodology and writing - revisions.

## Appendix

### Hierarchical Bayesian modeling

#### Model specification

Figure A1 shows a graphical illustration of the hierarchical Bayesian model we fit to empirical force matching data. In brief, for each participant and force level, we fit two parameters (mean and SD) to account for matching forces in the indirect condition. These participant-level parameters were samples from normal and gamma hyper-distributions, respectively, which were unique to each force level but common to all participants. For each participant, we fit two further parameters to account for matching forces in the direct condition across all force levels: these were the mean and SD of the attenuation factor, the key parameters of interest for our analysis. These participant-level parameters were again sampled from normal and gamma hyper-distributions common to all participants. Each parameter associated with a hyper-distribution (nodes outside the participant plate) was constrained by a hyperprior distribution. Below, we provide a full description of the model.

#### Indirect matching forces

We simultaneously fit the data from all conditions within each experiment. Our aim for the indirect reproduction conditions (*I*) was to measure the matching forces rather than to predict them, so we modelled the matching forces on these trials as samples drawn from a normal distribution, with a unique pair of mean and SD parameters for each target force level *t* and participant:

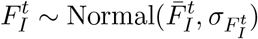

Note that for readability we suppress the participant index on all variables. For each participant, we set individual-level priors for the mean and standard deviation within each force level:

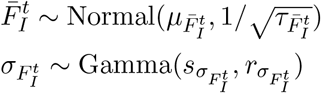

These individual-level parameters were, in turn, dependent on population-level hyperparameters, with hyperpriors:

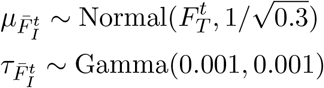

The shape (*s*) and rate (*r*) parameters of the gamma prior distribution on 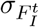 had priors defined in terms of the central tendency and width of the distribution:

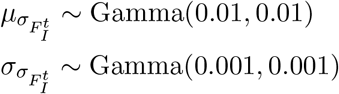

**Figure A1:**
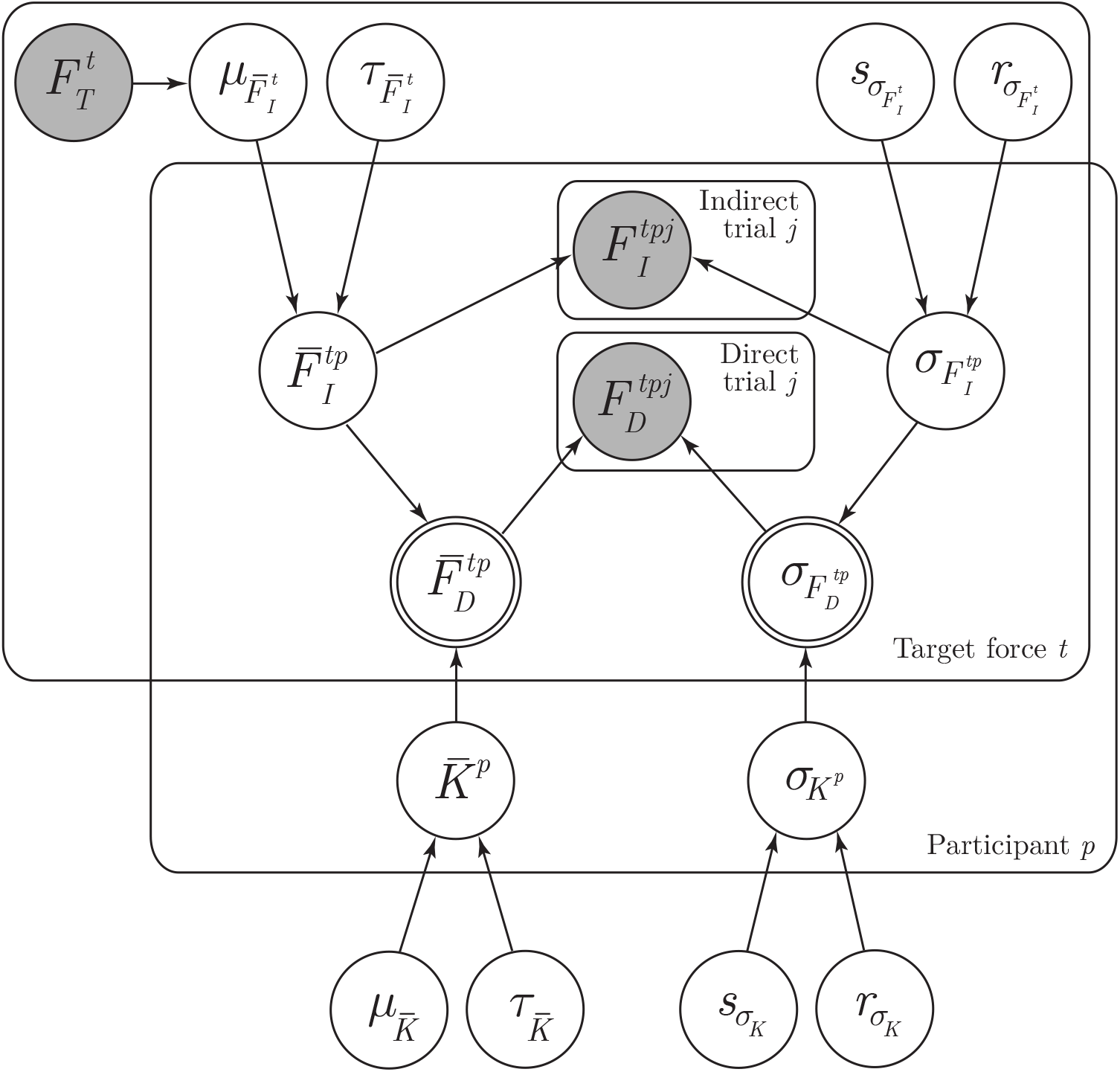
Graph describing the Bayesian hierarchical model. Unshaded and shaded circular nodes indicate unobserved and observed variables, respectively. Single-bordered nodes denote stochastic variables, while double-bordered nodes denote deterministic variables (as the mean direct matching force and standard deviation are fully determined by the stochastic means and standard deviations of the indirect force and attenuation factor). Plates indicate repetitive structures within the model. The additive version of the model is shown: the divisive model has an additional deterministic node corresponding to skewness of direct matching forces.

We then used a simple reparametrization to convert the mean and standard deviation into the shape and rate parameters [35]:

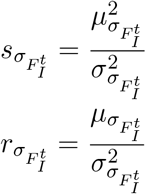

Parameterizing a gamma distribution in terms of the mean and standard deviation, as done here, results in a much more intuitive interpretation of the prior and resulting posterior distributions than when using shape and rate parameters. However, gamma distributions are typically positively skewed, which can affect their means due to their asymmetric tails. In this case, a potentially better measure of central tendency is the median. Because the median of a gamma distribution has no simple closed form, we opted to define priors over the mean instead. This approach offers a pragmatic compromise, particularly when using weakly informative priors. After fitting the model and obtaining the posterior distributions, we estimated the median numerically by first generating 10^7^ random samples using the accepted parameters on each MCMC iteration, and then calculating the median of those samples to estimate the posterior distribution of standard deviation of the attenuation. For completeness, we additionally computed the mode using the fitted shape and rate parameters of the gamma distribution [35]. Comparisons conducted on the mode estimates were largely consistent with those performed on the median estimates.

#### Direct matching forces

##### Subtractive model

In the subtractive model, direct matching forces are considered samples from a normal distribution with mean and SD determined by combining the indirect force matching parameters corresponding to the same target force 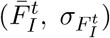 with the attenuation factor parameters 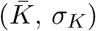, according to Eqs. 1 & 2:

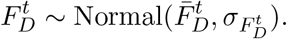

We set individual-level priors for the attenuation parameters:

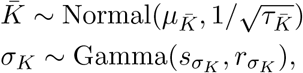

which are again dependent on population-level hyperparameters, with the following hyperpriors:

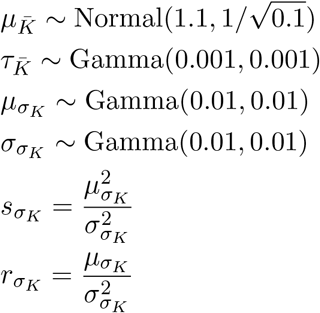

##### Divisive model

In the divisive model, attenuated output force on an individual trial is considered a sample from a skewed-normal distribution (see below) with mean, SD and skewness defined as in Eqs. 3–5:

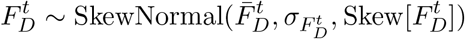

Individual- and population-level priors for the attenuation parameters and hyperparameters were set identically to the subtractive model (above) except for the population mean hyperprior for 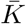 which was set to:

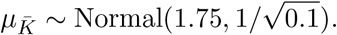

To illustrate the fitted parameters of the divisive model, we plot the mean population-level attenuation factor 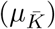 in Figure 5A. Following the described reparametrization of the Gamma distribution, we display the median population-level standard deviation of the attenuation factor 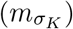 in Figure 5B.

### Results

#### Linear regression

**Figure A2:**
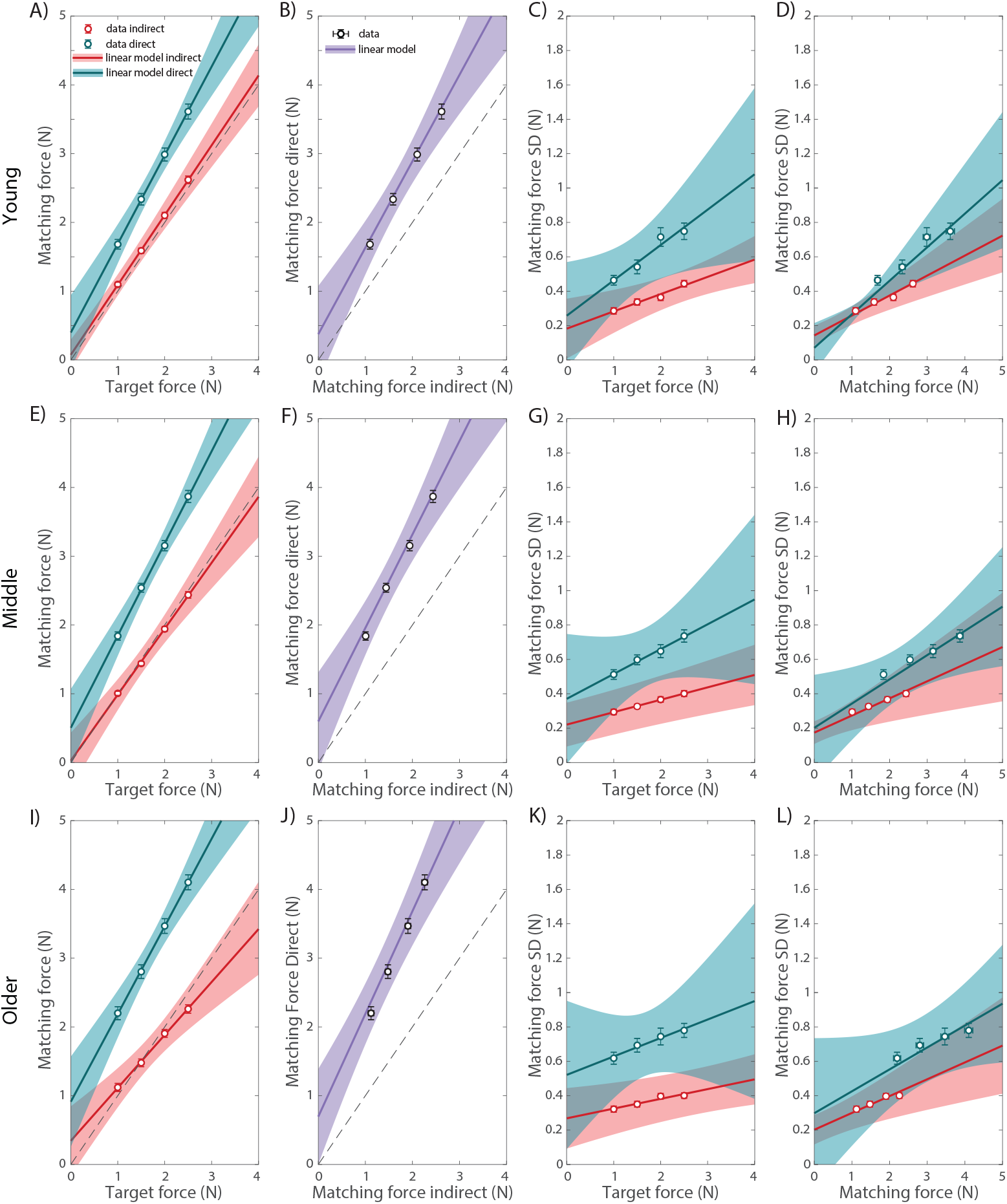
Matching forces and trial-to-trial variability in matching force in Study 2. (A) Mean matching force as a function of target force, for direct (cyan) and indirect (red) conditions. Data are shown as symbols and linear fits as lines. (B) Matching force in the direct condition plotted against matching force in the indirect condition, with linear fit. Symbols indicate mean matching forces for each target force level. (C) Standard deviation of matching force as a function of target force, for direct (cyan) and indirect (red) conditions, with linear fits. (D) Standard deviation of matching force as a function of mean matching force, with linear fits. Each symbol corresponds to a single target force level. (A-D) Young subjects. (E-H) Same as panels (A-D), but for middle-aged subjects. (I-L) Same as panels (A-D), but for older subjects. Error bars indicate *±*1SE, and shaded areas indicate 95% CI.

**Figure A3:**
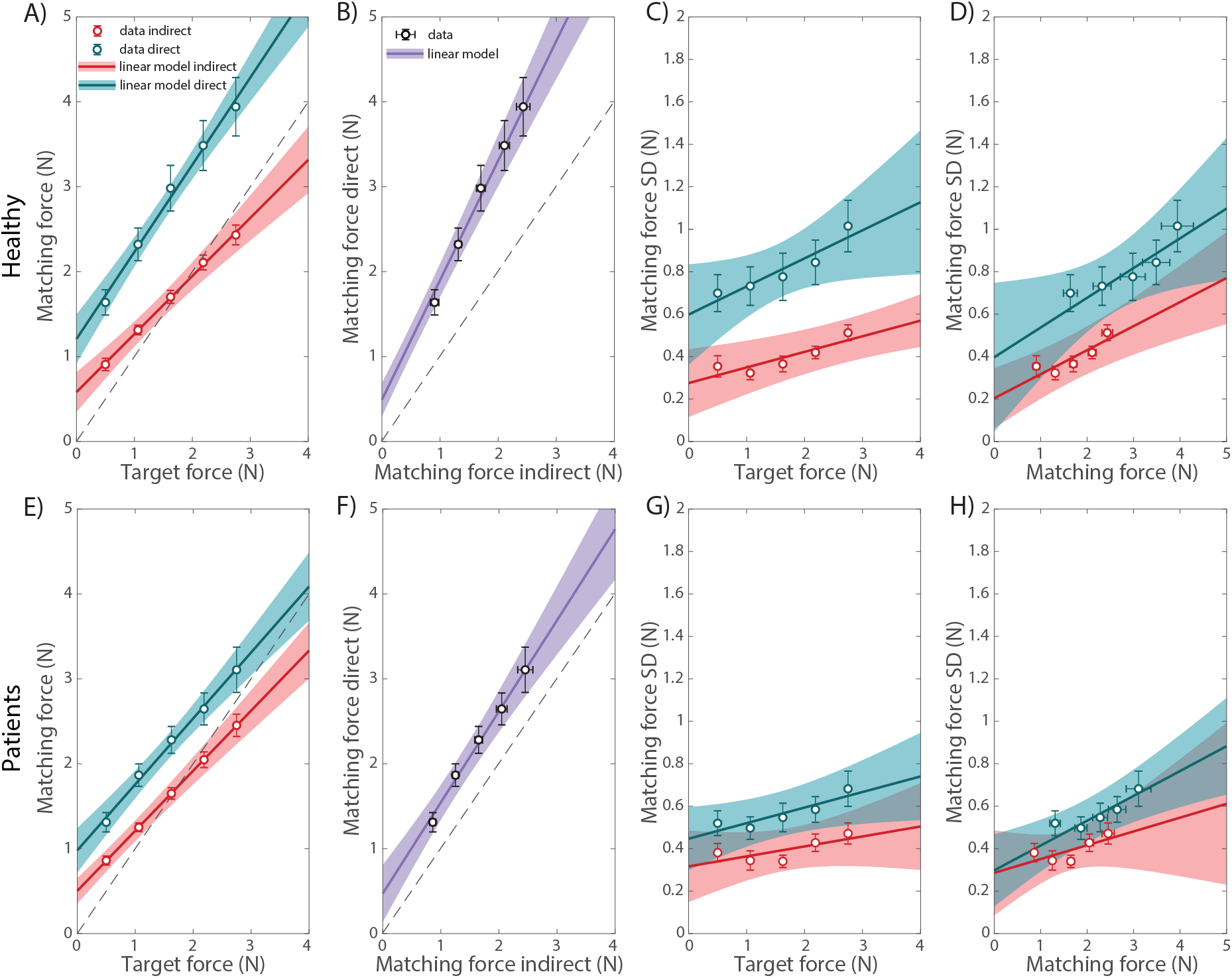
Matching forces and trial-to-trial variability in matching force in Study 3. (A) Mean matching force as a function of target force, for direct (cyan) and indirect (red) conditions. Data are shown as symbols and linear fits as lines. (B) Matching force in the direct condition plotted against matching force in the indirect condition, with linear fit. Symbols indicate mean matching forces for each target force level. (C) Standard deviation of matching force as a function of target force, for direct (cyan) and indirect (red) conditions, with linear fits. (D) Standard deviation of matching force as a function of mean matching force, with linear fits. Each symbol corresponds to a single target force level. (A-D) Healthy subjects. (E-H) Same as panels (A-D), but for patients subjects. Error bars indicate *±*1SE, and shaded areas indicate 95% CI.

#### Divisive model

**Figure A4:**
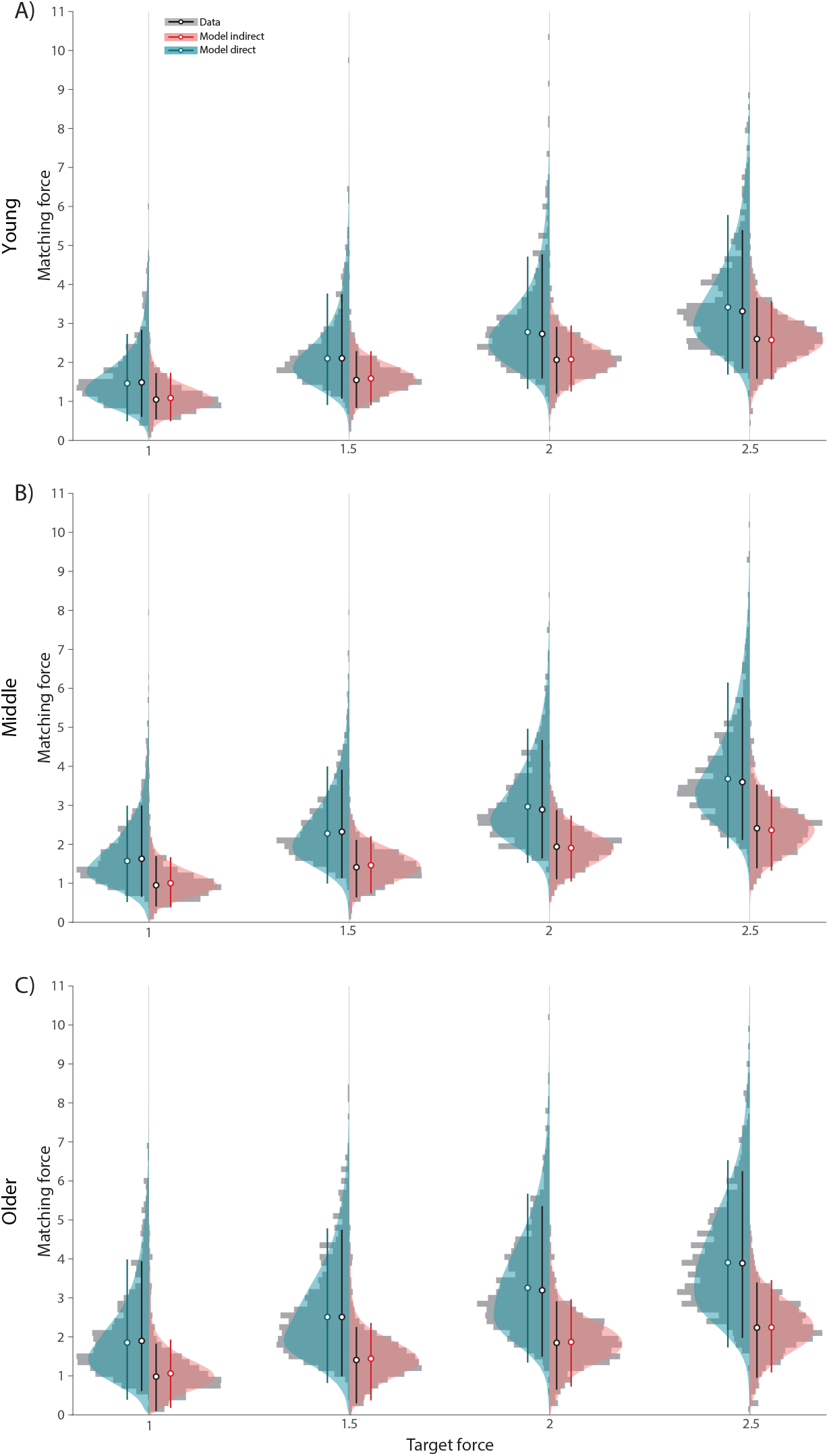
Posterior predictive densities for Study 2. (A) Young subject. (B) Middle-aged subjects. (C) Older subjects. Grey histograms show matching forces pooled across all participants. Coloured distributions show simulated matching forces based on posterior predictive density. The circles with error bars show the median and 90% HDI for data (black) and simulated trials for the same condition (cyan: direct, red: indirect).

**Figure A5:**
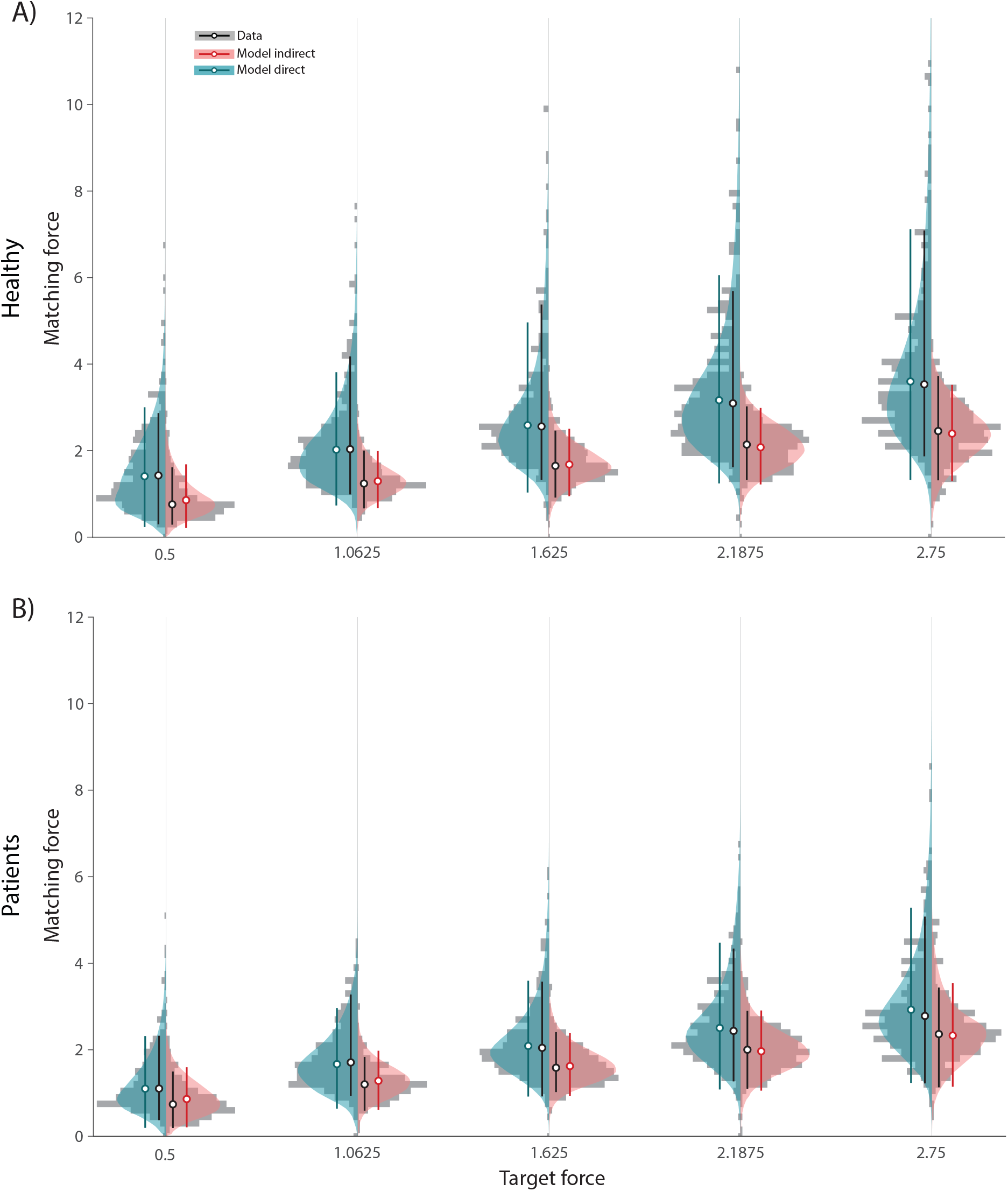
Posterior predictive densities for Study 3. (A) Healthy subject. (B) Patients subjects. Grey histograms show matching forces pooled across all participants. Coloured distributions show simulated matching forces based on posterior predictive density. The circles with error bars show the median and 90% HDI for data (black) and simulated trials for the same condition (cyan: direct, red: indirect).

#### Subtractive model

**Figure A6:**
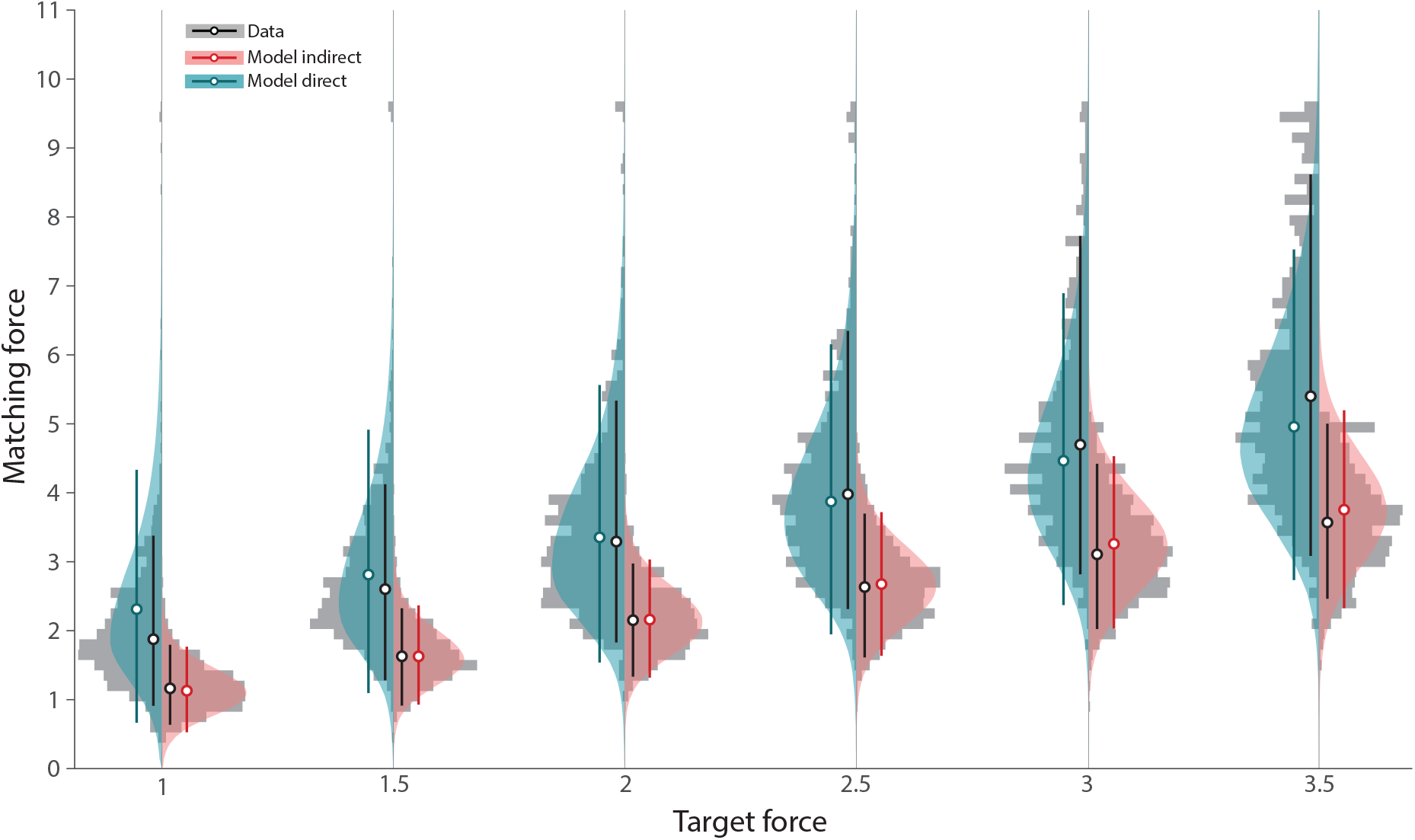
Subtractive model posterior predictive densities for Study 1. Grey histograms show matching forces pooled across all participants. Coloured distributions show simulated matching forces based on posterior predictive density. The circles with error bars show the median and 90% HDI for data (black) and simulated trials for the same condition (cyan: direct, red: indirect).

**Figure A7:**
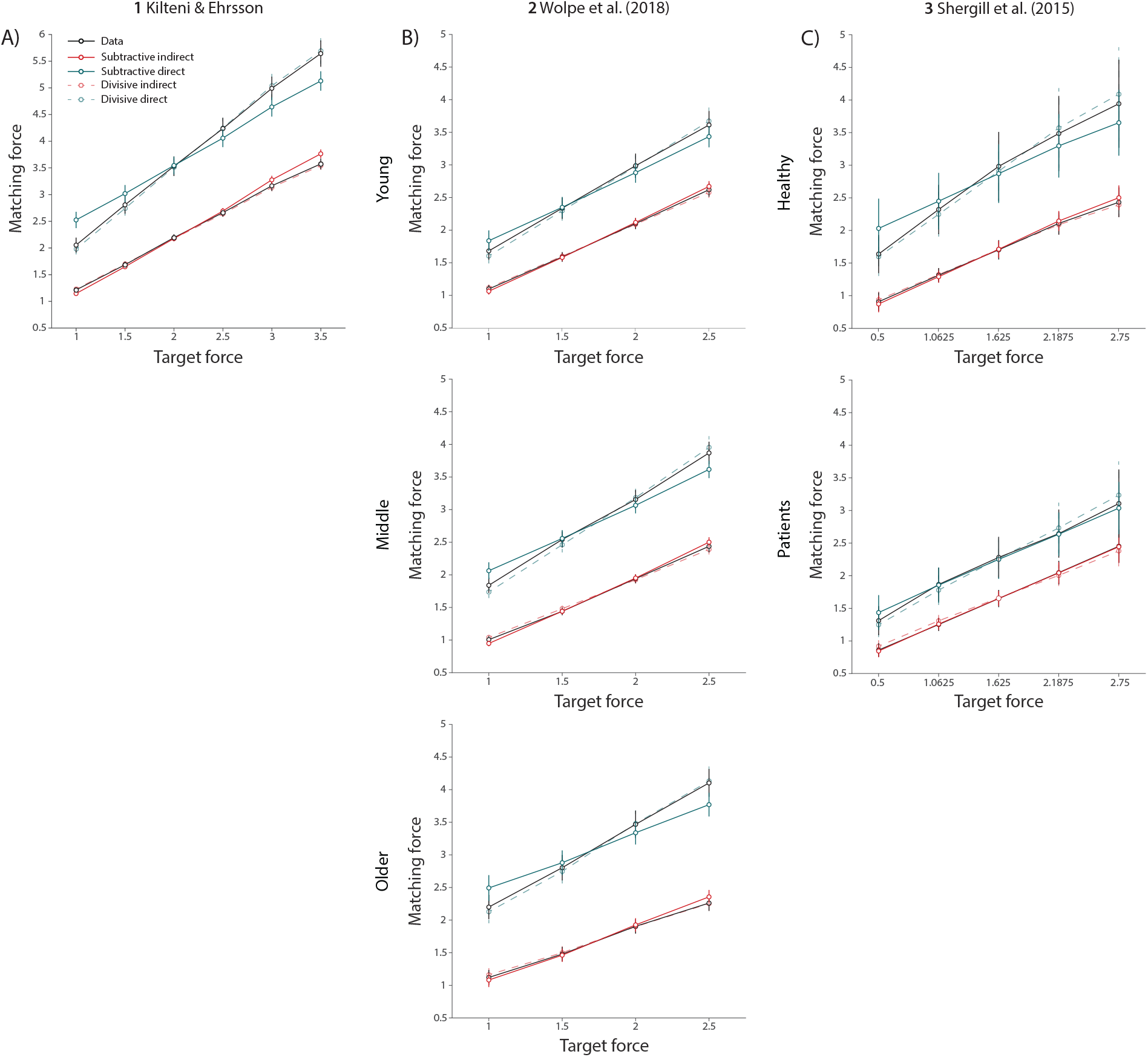
Subtractive model posterior predictive check. (A) Study 1. (B) Study 2 (from top to bottom: young, middle, and older subjects); (C) Study 3 (from top to bottom: healthy and patient subjects). The solid and dashed lines show fits of the subtractive and divisive model, respectively. The circles with error bars show the mean and 95% CI.

**Table A1:** Summary of Subtractive model

| no | Sample | Attenuation factor | | SD of attenuation factor | | Bayesian $p$-value |
| --- | --- | --- | --- | --- | --- | --- |
|  |  | Median | 90% HDI | Median | 90% HDI |  |
| 1 | Study 1 | 1.36 | [1.24, 1.49] | 0.726 | [0.657, 0.803] | $0.34 \leq \text{all } p \leq 0.61$ |
| 2 | Study 2 young | 0.76 | [0.62, 0.90] | 0.48 | [0.403, 0.569] | $0.43 \leq \text{all } p \leq 0.54$ |
| 3 | Study 2 middle | 1.11 | [1.00, 1.23] | 0.526 | [0.467, 0.586] | $0.40 \leq \text{all } p \leq 0.57$ |
| 4 | Study 2 older | 1.41 | [1.26, 1.55] | 0.646 | [0.567, 0.726] | $0.39 \leq \text{all } p \leq 0.60$ |
| 5 | Study 3 healthy | 1.15 | [0.77, 1.53] | 0.727 | [0.520, 0.955] | $0.39 \leq \text{all } p \leq 0.55$ |
| 6 | Study 3 patients | 0.60 | [0.36, 0.84] | 0.393 | [0.261, 0.528] | $0.46 \leq \text{all } p \leq 0.52$ |

### Skew-normal approximation to the product of normals

If *X* ∼ *N* (*µ*_*x*_, *σ*_*x*_) and *Y* ∼ *N* (*µ*_*y*_, *σ*_*y*_), for *Z* = *XY*,

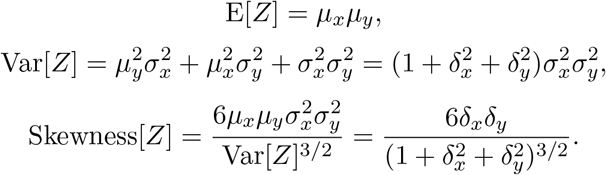

where 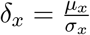 and 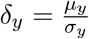.

This distribution can be approximated by a skew-normal distribution with the same moments. The skew-normal has pdf,

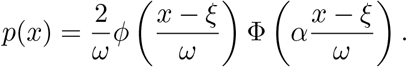

Defining

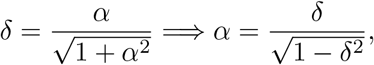

for −1 *< δ <* 1.

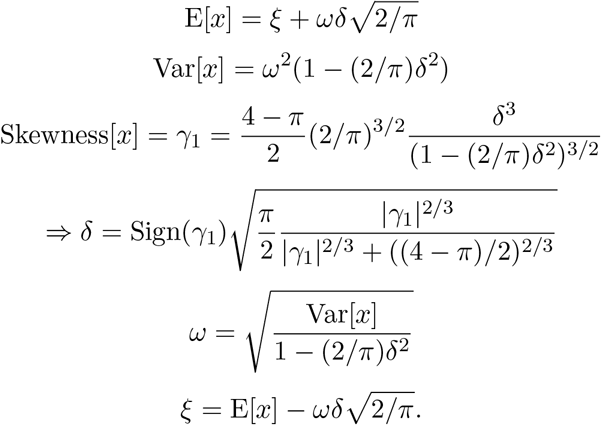

